# Current and potential contributions of the Gulf of Lion Fisheries Restricted Area to fisheries sustainability in the NW Mediterranean Sea

**DOI:** 10.1101/2020.02.22.960914

**Authors:** Daniel Vilas, Marta Coll, Xavier Corrales, Jeroen Steenbeek, Chiara Piroddi, Diego Macias, Alessandro Ligas, Paolo Sartor, Joachim Claudet

## Abstract

Many commercial species of the world are overexploited resulting in substantial reductions of biomass and ecological changes. Spatial-temporal restrictions of fishing activities are important measures used for the management of marine stocks. However, evidence of whether fishing bans benefit whole ecosystems is still scant. Here, we developed a food-web model approach using the Ecopath with Ecosim (EwE) model representing the Fisheries Restricted Area (FRA) of the Gulf of Lion ecosystem (CoSEGoL model) prior to the establishment of the fisheries restrictions (2006-2008) to characterize the structure and functioning of the ecosystem before and after its establishment. The constructed food-web model was, then, fitted to available time series of data from 2008 to 2016 to verify whether this FRA has contributed to recovery of target demersal species and the demersal community. The fitted model was used to explore alternative future management scenarios to explore feasible management options in order to ensure a full ecosystem recovery under climate change conditions. Both small positive and negative ecosystem changes occurred between prior and after the establishment of the FRA, potentially revealing a lack of protection efficiency and/or enforcement. Scenarios of management options under plausible climate futures revealed possible recovery of targeted species, especially European hake. The study highlighted the importance of considering trophic interactions between predators and prey to identify trade-offs and synergies in fisheries management outcomes and the need to consider both fishing and climate dynamics.

## 1. Introduction

Fishing is considered one of the most harmful stressors of marine ecosystems [1], with impacts on habitat [2], biodiversity [3] and ecosystem structure [4]. Overexploitation of marine resources is widely distributed [5], and it has substantially reduced fish biomass and caused significant ecological changes in the global ocean [6,7]. In the Mediterranean Sea, many assessed demersal stocks are either fully exploited or overexploited [8]. This situation is not substantially improving as exploitation rate is increasing [8,9], and selectivity is decreasing [8,10].

Spatial-temporal restrictions of fishing activities and the establishment of technical measures are the main management tools used in the Mediterranean Sea for marine exploited stocks [9]. Under the European Union (EU) and the General Fisheries Commission for the Mediterranean (GFCM), legal frameworks concerning the establishment of spatial-temporal restrictions of fishing activities come mostly from two regulations [11,12] and one recommendation [13]. The first regulation indicates the definition for fishing protected areas and spatial-temporal restrictions in the Mediterranean Sea, while the second one establishes the need to advance the spatial-temporal measures for recovering populations of demersal stocks in the Western Mediterranean Sea. The recommendation states that the use of trawl nets in waters deeper than 1000 metres shall be prohibited to protect little-known deep-sea benthic habitats in the Mediterranean. Waters below 1000 meters were officially declared as a Fisheries Restricted Area (FRA) by the EU Commission in 2016 [8]. In addition, since 2006, eight FRAs have been established to ensure the protection of deep-sea sensitive habitats and essential fish habitats in well-defined areas of the Mediterranean Sea. Among these is the continental slope of the Eastern Gulf of Lion (CoSEGoL) FRA, the only FRA located in the Western Mediterranean Sea outside territorial waters. The CoSeGoL FRA was established in 2009, following a Recommendation by GFCM (GFCM/33/2009/1) [14], which froze the fishing effort in the area, pending the delivery of additional information by the Scientific Advisory Committee on Fisheries (SAC). In fact, the SAC had advised “to ban the use of towed and fixed gear and longlines for demersal resource in an area of the continental shelf and slope of the eastern Gulf of Lion” [14].

A lack of effective management and/or enforcement in restricted areas to fishing has been identified as a major flaw in the Mediterranean Sea management system [15], and it is likely that the FRAs are not reaching their total effectiveness due to increasing illegal or unreported fishing activities inside these areas or due to insufficient measures. For example, Petza *et al.*, [16] reviewed the effectiveness of several national FRAs in the Aegean sea and found that more than 50% of the studied FRAs (n=516) were slightly effective because of their multi-criteria based on the management of FRAs.

By 2020 the 10% of the Mediterranean Sea should be protected [17] to ensure the improvement of the status of fish stocks and fisheries. Well established and effective FRAs could contribute to increase the protected surface in the Mediterranean Sea [18,19]. Currently, official spatial protection in the Mediterranean Sea covers more than 10% of its surface [20], although most of these areas are poorly protected or unprotected [21] and the surface of fully protected areas is around 0.04% [22].

In such a context, it is important to assess whether the proposed FRAs are effectively delivering the benefits they are expected to. Here, we developed a food-web model to evaluate the effectiveness of the CoSEGoL FRA to rebuild and protect demersal commercial stocks in the North Western Mediterranean Sea and to ensure a resilient structure and functioning of the ecosystems. Specifically, we characterized the structure and functioning of the area; we assessed whether the FRA helped recover targeted demersal species and the demersal community since its establishment; and we explored the viability of alternative management options under climate change. To this end we developed a food web model using the Ecopath with Ecosim (EwE) approach [23,24] representing the CoSEGoL area (2006-2008) prior to the establishment of the FRA. The food-web model was fitted to available time series of data from 2008 to 2016 using the temporal dynamic module Ecosim [24,25] to simulate how the structural and functional traits of the ecosystem changed since the establishment of the FRA, and to verify if its establishment resulted in the recovery of commercially targeted species. The fitted model was then used to explore alternative future management scenarios under climate change conditions (accounting for changes in the water temperature and primary productivity dynamics), following similar approaches applied in other modelling studies [26–28]. This study complements existing modelling studies of protected areas in the Mediterranean Sea [29–32] using the EwE approach, by explicitly representing the FRA in the basin..

## 2. Material & Methods

### 2.1. Study area

The CoSEGoL FRA is located in the Gulf of Lion in the Northwestern Mediterranean Sea, bounded by the following geographic coordinates: 42°40’N, 4°20’ E; 42°40’N, 5°00’ E; 43°00’N, 4°20’ E; 43°00’N, 5°00’ E (Figure 1). The Gulf of Lion is one of the most productive regions of the Mediterranean Sea because of the inputs from the Rhone river and experiences annual upwelling [33]. The bathymetry of the CoSEGoL FRA ranges from 100 to 1500 meters and covers an area of 2,051 km^2^ [34]. This area has been identified as containing essential fish habitats (nurseries and spawning areas) for European hake (*Merluccius merluccius*) and other commercials species [35]. It is characterized by an intricate network of submarine canyons [36], and important benthonic communities of echinoderms, gorgonians, sponges [37] and deep-sea corals [38,39] occur in the area.

**Figure 1.**
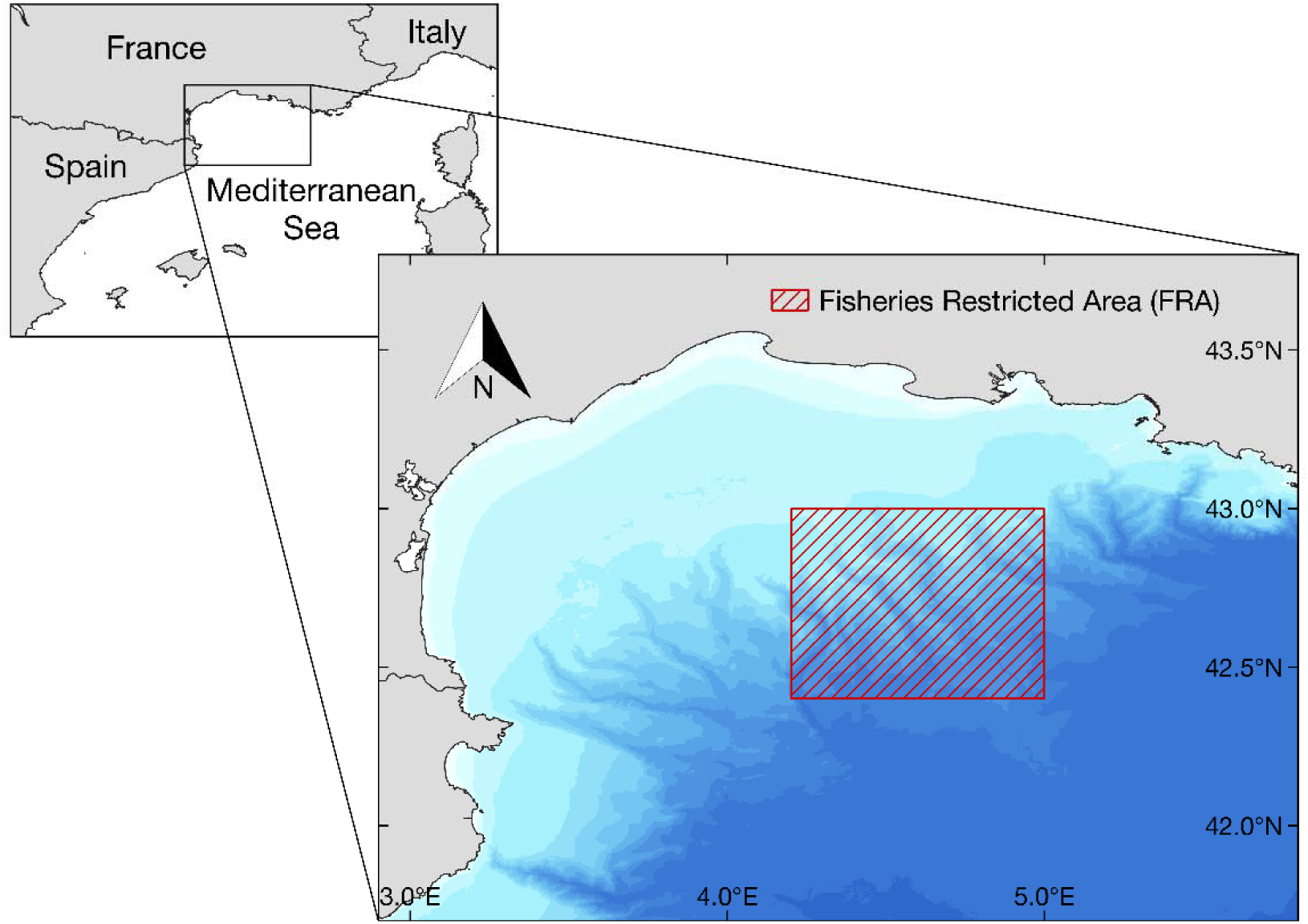
Location of the continental slope of the Eastern Gulf of Lion Fisheries Restricted Area (Northwestern Mediterranean Sea).

The CoSEGoL FRA is historically exploited by Spanish bottom trawlers (BTW), Spanish longliners (LON) and French midwater trawlers (MTW) [14]. French trawlers are the main component of the fleet exploiting the marine resources of the Gulf of Lion, and can be divided in two main components: one directed to the catch of small pelagic fish, and the other exploiting a great diversity of demersal species [37]. The aim of CoSEGoL FRA was to protect very important spawning stocks of several species of fishes of importance in the Northwestern Mediterranean fisheries, the most important one being European hake, and also including anglerfish (*Lophius piscatorius*), Norway lobster (*Nephrops norvergicus)* and the blue and red shrimp (*Aristeus antennatus*), while conserving accompanying species (blue whiting *Micromesisitius poutassou* and silver sabbardfish *Lepidopus caudatus*) [37].

In the recommendation from which the CoSEGoL FRA was adopted [14], the GFCM called for ensuring that fishing effort for demersal stocks of vessels using towed nets, bottom and mid-water longlines, and bottom-set nets shall not exceed the level of fishing effort applied in 2008. Officially, the fleet operating that area in 2008 was composed by 29 fishing vessels, 70% from France and 30% from Spain [14].

### 2.2. Ecosystem modelling approach

The CoSEGoL FRA model was developed using the Ecopath with Ecosim ecosystem modelling approach (EwE, version 6.6) and it was built using the best available information to represent the FRA ecosystem just before its establishment. Specifically, the model represented a situation of the CoSEGoL FRA for 2006-2008 time period. Subsequently, an Ecosim model representing the CoSEGoL FRA ecosystem during the 2008−2016 period was fitted to time series of historical data. (See detailed information about ecosystem modelling approach in Supplementary material Appendix A).

### 2.3. Model parametrization

The CoSEGoL FRA model represented the state of the ecosystem in 2006-2008, previously to the official establishment of the FRA in 2009. Information about species presence and their biomasses was aggregated into functional groups (FGs) of species or groups of species clustered according to key information about their trophic ecology, commercial value, and abundance in the ecosystem. We used the same meta-web structure as defined for the SafeNet Project^1^ Western Mediterranean Sea model [43]. We adapted this meta-web structure to local conditions by removing those FGs that did not occur in the study area. The final food-web structure of the CoSEGoL FRA model contains 72 functional groups (five marine mammals, one seabird, one sea turtle, 13 pelagic fishes, 24 demersal fishes, four cephalopods, 18 invertebrates, two zooplankton, two phytoplankton and two detritus groups) (Supplementary material Table B.1).

FGs’ biomasses were obtained from different sources from the study area or surrounding areas (see Supplementary Material Table B.1 and C.1. for details on the parameterization of each functional group). Most of the biomasses of demersal and benthic species were calculated from the EU-funded Mediterranean International bottom Trawl Surveys project [44], carried out from spring to early summer (April to June) from 1994 to the present. Species biomass was estimated for each haul as the total weight of each species (kg) per km^2^ of trawling. This information was extracted from the MEDITS dataset to account for bathymetric sampling per strata. For pelagic species, we also used the data available from the EU-funded Mediterranean International Acoustic Survey (MEDIAS), which contained information of abundance and biomass per Geographical Sub-Area (GSA).

Annual production (P/B) and consumption (Q/B) rates were either estimated using empirical equations [45], or taken from literature or from other models developed in the Mediterranean Sea [43] (Supplementary Material Table B.1 and C.1.). The diet information was compiled using published studies (Supplementary Material Table B.1) on stomach content analyses, giving preference to local or surrounding areas (Supplementary Material Table C.2). We used the Diet Calculator (Steenbeek 2018), a custom-built extraction tool that facilitates the process of vetting and incorporating diet data into EwE. Drawing on a large library of published diet studies, the Diet Calculator selects the most likely suitable diet studies for a specific model area, based on a weighted evaluation of diet study characteristics, and generates a diet composition matrix with accompanying pedigree index for each predatory functional group. For migratory species (large pelagic fishes, sea birds, turtles and dolphins), we set a fraction of the diet composition as import based on the time that these species feed outside the system [23,45].

Fisheries data were obtained from different sources (database, literature and unpublished data) (Supplementary Material Table B.1. and C.1.). Available fishery data were not geolocated, and so we had to scale catches by the fishing area where operates each fleet. We divided fisheries in three commercial fishing fleets for the CoSEGoL FRA model (Spanish Bottom trawlers, Spanish longliners and French Midwater trawlers). We calculated catches in two different ways: 1) for French fleets, we scaled total catches [46] by FRA area belonging to Gulf of Lion area, and then by the number of vessels working in the study area [14]; and 2) for Spanish fleets, we obtained landings from official dataset of the Regional Government of Catalonia managed by the Institute of Marine Sciences (ICM-CSIC) [47], and were scaled to the area where these fleet were operating.

### 2.4. Quality of the model

The quality of the models were evaluated using the EwE pedigree routine, which allows assigning a measure of confidence to key input parameters (B, P/B, Q/B, diets and catches) [23,24]. All pedigree values were manually calculated except for diets, which were obtained from the Diet Calculator algorithm [49]. The algorithm computes a total pedigree value for each diet record as a weighted average of four attributes assigned to each diet study (region and year of collection, data representativeness of the species population, and data collection method). Pedigree values were used to identify parameters with low quality that could be modified during the balancing procedure, and were used to calculate the pedigree index, which varies between zero (lowest quality) and one (highest quality) [24], for the FRA model.

### 2.5. Fitting to time series procedure

Relative fishing effort data available for the fishing fleets included in the model were used to drive the model. Due to the lack of local fishing effort time series, and to test the hypothesis of compliance and enforcement failure in the CoSEGoL FRA, we tested alternative relative fishing effort time series that considered annual declines (−1%, −5%, - 10%), annual increases (+1%, +5%, +10%) or no changes in effort with time. These changes were applied to all fisheries in the model. Available absolute or relative observed biomass time series were incorporated in Ecosim to compare the model outputs to observations.

### 2.6. Model analyses and ecological indicators

The food web structure of the CoSEGoL FRA ecosystem before and after the establishment of the FRA was visualized using a flow diagram built from the biomass and TL (output) of each FG, and the direct trophic interactions among them. The TL identifies the position of organisms within food webs by tracking the source of energy for each organism, and it is calculated by assigning primary producers and detritus a TL of 1 (e.g. phytoplankton), and consumers to a TL of 1, plus the average TL of their prey weighted by their proportion in weight in the predator’s diet [52].

With both before and after FRA models, the mixed trophic impact (MTI) analysis was performed to quantify direct and indirect trophic interactions among functional groups [53]. This analysis quantifies the direct and indirect impacts that a hypothetical increase in the biomass of one functional group would have on the biomasses of all the other functional groups in the ecosystem, including the fishing fleets. We also used Valls keystone index [54] to identify keystone species in both before and after FRA models. A keystone species is a predator species that shows relatively low biomass but has a relatively important role in the ecosystem [55].

Several additional ecological indicators were computed to describe the state and functioning trend of the CoSEGoL FRA before and after the establishment of the fisheries restrictions following [56]:

#### Biomass-based

These indicators are calculated from the biomass of components included in the food-web model. We included five biomass-based indicators: biomass of demersal species (t·km^−2^·year^−1^) biomass of fish species (t·km^−2^·year^−1^), biomass of commercial species (t·km^−2^·year^−1^), biomass of predatory species (t·km^−2^·year^−1^) and biomass of invertebrates species (t·km^−2^·year^−1^).

#### Trophic-based

These indicators reflect the TL position of different groups of the food web. Trophic level indicators may reflect ecosystem “health” because fishing pressure removing predators can cause a decline in the trophic level of the catch and/or the community [52]. We selected four trophic-based indicators: TL of the community (TLc), TL of the community including organisms with TL ≥ 2 (TL2), TL of the community including organisms with TL ≥ 3.25 (TL3.25) and TL of the community including organisms with TL ≥ 4 (TL4).

#### Flows-based

We used two indicators related to total flows of the system. The Average Path Length (APL, μ) is defined as the average number of groups that flows passes through and is an indicator of stress [57]. The Finn’s Cycling Index (FCI, %) is the fraction of the ecosystem’s throughput that is recycled [58].

#### Catch-based

These indicators are based on catch and discard species data. We included six indicators: total catch (t·km^−2^·year^−1^), total demersal catch (t·km^−2^·year^−1^), total fish catch (t·km^−2^·year^−1^), total invertebrates catch (t·km^−2^·year^−1^), total discarded catch (t·km^−2^·year^−1^) and trophic level of the catch.

### 2.7. Assessment of FRA impact and uncertainty

After fitting the model to time series using Ecosim, we investigated if the establishment of the CoSEGoL FRA resulted in noticeable changes in the structure and functioning of the ecosystem. We compared the ecosystem structure and functioning before and after the establishment of the FRA using the baseline model (2008) and a second FRA model that was extracted at the end of the fitting time period (2016). FRA effectiveness and compliance was measured through changes in ecological and keystone species indicators to discern expected biomass increases according to theory [59]. For example, a positive trend in the biomass of a targeted species is to be expected in a FRA after several years of its protection [18]. In addition, changes in mixed trophic impacts (MTI) from the industrial fleets were examined to quantify the direct and indirect impact of each fleet on functional groups, their potential competitions and trade-offs.

Pedigree and associated confidence intervals for key input values were used in the EwE Monte Carlo (MC) routine to evaluate input parameter uncertainty over time (Supplementary Material Table D.1) [24,45]. 200 MC simulations were run, and 95% and 5% percentile confidence intervals (CIs) were calculated for main target species biomasses and ecological indicators focussing on *M. merluccius, L. piscatorius, N. norvergicus, A. antennatus, M. poutassou* and *L. caudatus*. The significance and correlation between our suite of ecological indicators and time were measured using the non-parametric Spearman rank correlation coefficient [60]. To evaluate the impact of the CoSEGoL FRA on the fisheries, catch-based indicator trends were examined over time to capture changes of the potential effects of the FRA establishment. This procedure to capture uncertainty was developed to evaluate historical changes (2008-2016) and the forecasting scenarios (see section below).

### 2.8. Future alternative management simulations

After the model was fitted to data from 2008 to 2016, eight future scenarios (Table 1) were tested in order to evaluate future alternative management scenarios and their potential effects on marine resources and the ecosystem structure and functioning in the 2017-2040 period. The original configuration of the dynamic model was used as a baseline simulation keeping parameters with default values from 2017 to 2040 (Business as usual - BAU). The rest of the scenarios applied new fishing regulations: for instance, scenario “50” simulated a decreasing 50% of fishing effort, scenario “100” simulated a decreasing 100% of fishing effort, and scenario “F_msy_” simulated fishing at Maximum Sustainable Yield (F_msy_) in comparison to fishing at F current (using fishing mortality levels of 2016). To obtain more realistic predictions, we considered future projections of temperature and primary production change.

**Table 1.** List of fisheries and RCP scenarios.

| Scenario | Fishing regulation | Temperature | PP |
| --- | --- | --- | --- |
| BAU 4.5 | Kept at levels 2016 | RCP4.5 | RCP4.5 |
| BAU 8.5 | Kept at levels 2016 | RCP8.5 | RCP8.5 |
| 100 4.5 | Reducing 100% fishing effort | RCP4.5 | RCP4.5 |
| 100 8.5 | Reducing 100% fishing effort | RCP8.5 | RCP8.5 |
| 50 4.5 | Reducing 50% fishing effort | RCP4.5 | RCP4.5 |
| 50 8.5 | Reducing 50% fishing effort | RCP8.5 | RCP8.5 |
| F<sub>msy</sub> 4.5 | Maximum sustainable yield | RCP4.5 | RCP4.5 |
| F<sub>msy</sub> 8.5 | Maximum sustainable yield | RCP8.5 | RCP8.5 |

To obtain the F_msy_ values, we first reviewed fishing mortality values at current levels (F_current_) and F0.1 (defined as the fishing mortality at which the slope of the Yield per Recruit curve is 10 percent of its slope at the origin) reported by the GFCM and the Scientific, Technical and Economic Committee for Fisheries (STECF) in the last evaluations of Western Mediterranean marine resources (Supplementary material, Table D.2.). Values of F_current_ and F_msy_ for European sardine (*Sardina pilchardus*) and European anchovy (*Engraulis encrasicolus*) were obtained from EU Tender SPELMED for GSA06 and GSA07 [61]. We estimated the reduction of fishing mortality comparing F_current_ with F0.1 for evaluated species, which yielded an average reduction of 64% for the CoSEGoL FRA. This estimate was applied to the rest of the commercial species that were not assessed but also occurred in the model as fisheries targeted or by-catch species.

For the environmental variables (sea water temperature and primary production) (Supplementary material, Figures E.1. and E.2), we used projections of the Med-ERGOM hydro-dynamical biochemical model under two contrasting scenarios of greenhouse emissions (RCP4.5 and RCP8.5) [62,63] (Table 1). To consider changes in sea water temperature, we used the environmental response functions of Ecosim, which links the species or FGs dynamics to the environmental drivers. We first obtained the response functions from AquaMaps [64], which is a global database on species distribution. These environmental response functions are given as curves showing minimum and maximum tolerance levels and 10th and 90th preferable quintiles to the environmental parameters (in our case, temperature). The final environmental preferences for each FG were obtained by weighting the values of the species included in a FG to their relative biomass. Finally, selected ecological indicators and biomass predictions of targeted species were extracted in 2025 and 2040 and were used to assess the effects of future alternative simulations over time.

## 3. Results

### 3.1. Baseline parameterization, model quality and temporal fitting

The pedigree index of the CoSEGoL FRA model (0.50) revealed that input data was of acceptable quality when compared to the distribution of pedigree values in other existing models [65]. However, the pedigree value of the CoSEGoL FRA was lower than other published EwE models for the NW Mediterranean Sea [66,67].

The best fitted model was obtained for an annual increase in fishing effort of 5% (F_+5_) (Supplementary material Table F.1.). The parameterization with 30 vulnerabilities (trophic interactions between predators and their prey) and 6 spline points was identified as the best model based on the AIC test criteria (Supplementary material Table F.1). However, the best fitting model did not reproduce observed trends of some target species; these were obtained for a scenario with an increase of 10% in fishing effort. This model was adopted as most likely representative for the ecosystem because of its capability to best reproduce the trends in target species over time (Supplementary material Table F.1) with exception of Norway lobster (SS 11.87) - one of the target groups of the study (Figure 2).

**Figure 2.**
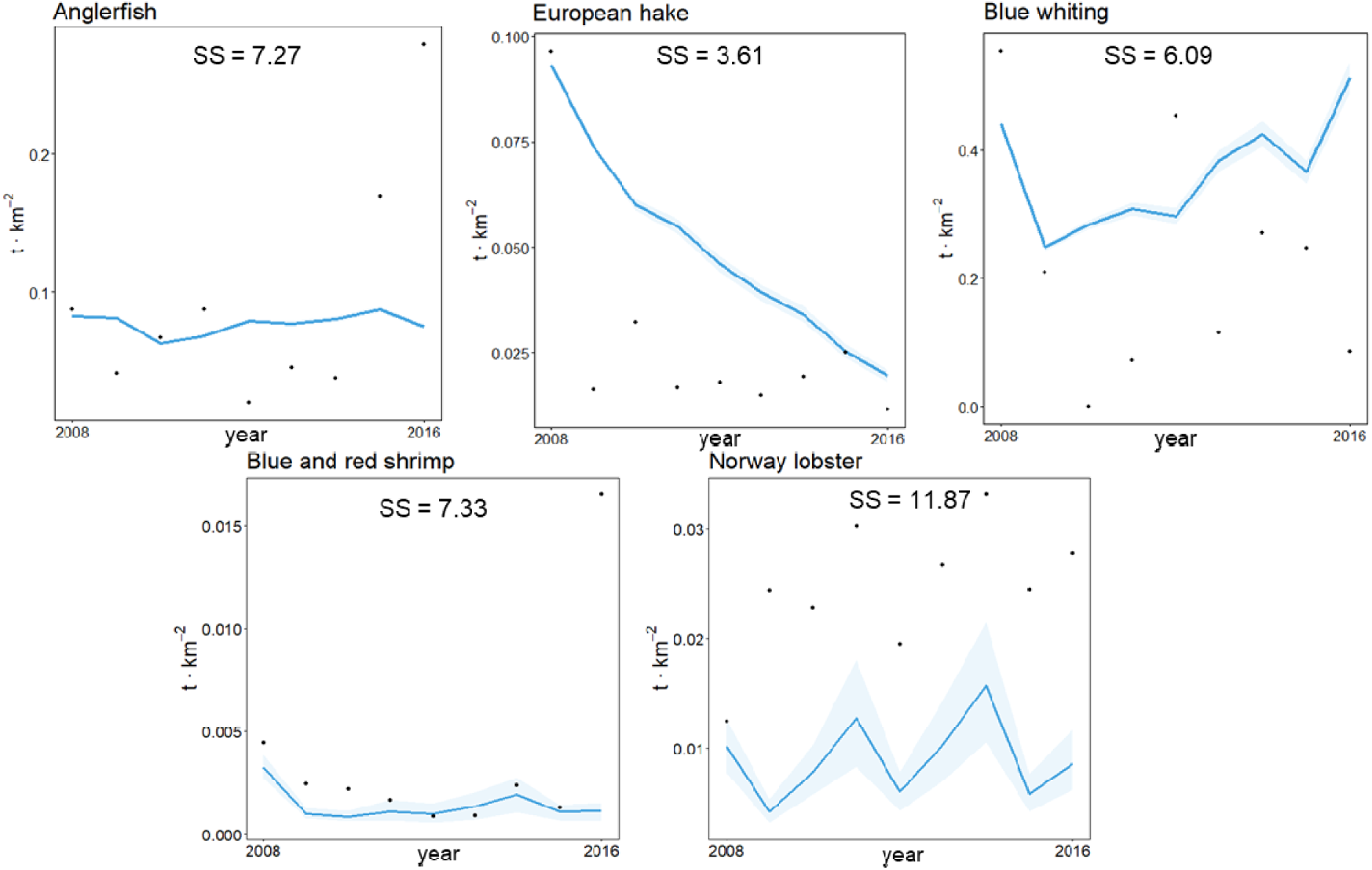
Predicted (solid lines) versus observed (dots) biomass (t·km^−2^) for targeted groups in the continental slope of the Eastern Gulf of Lion Fisheries Restricted Areaecosystem model for the period 2008-2016. Blue shadows represent the 5% and 95% percentiles obtained using the Monte Carlo routine. Sum of squares (SS) values indicate their contribution for the total SS.

### 3.2. Ecosystem structure and functioning

The structure and functioning of the ecosystem changed between prior and after the FRA establishment. The flow diagram showed higher trophic levels for the model prior the establishment of the FRA (Figure 3). Both models highlighted the same FGs for Valls keystone index (Figure 4), although keystone index values for individual groups differed from one ecosystem state to the other. Ecological indicators showed generally small variation. Of biomass-based indicators, only invertebrates showed noticeable differences (Figure 5), with an increase from 2008 to 2016. The TL community, TL community 2 and APL decreased after the implementation of the FRA (Figure 6 and 7).

**Figure 3.**
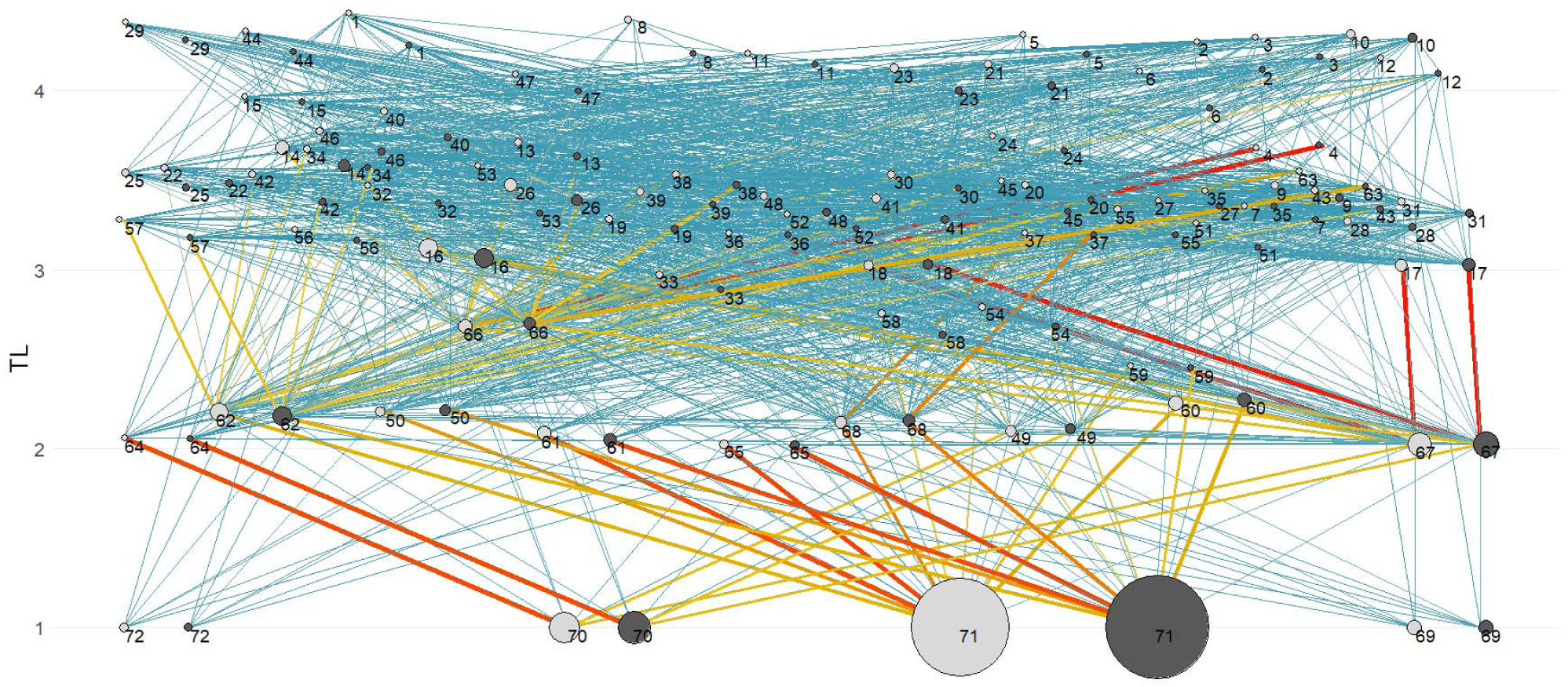
Flow diagram of the continental slope of the Eastern Gulf of Lion Fisheries Restricted Areaecosystem before its establishment (year 2008, light grey) and after (year 2016, dark grey). The size of each circle is proportional to the biomass of the functional group. The numbers identify the functional groups of both CoSEGoL FRA models (Table A.1). The width and color of trophic links indicate the magnitude of the trophic flows (low - blue; high - red).

**Figure 4.**
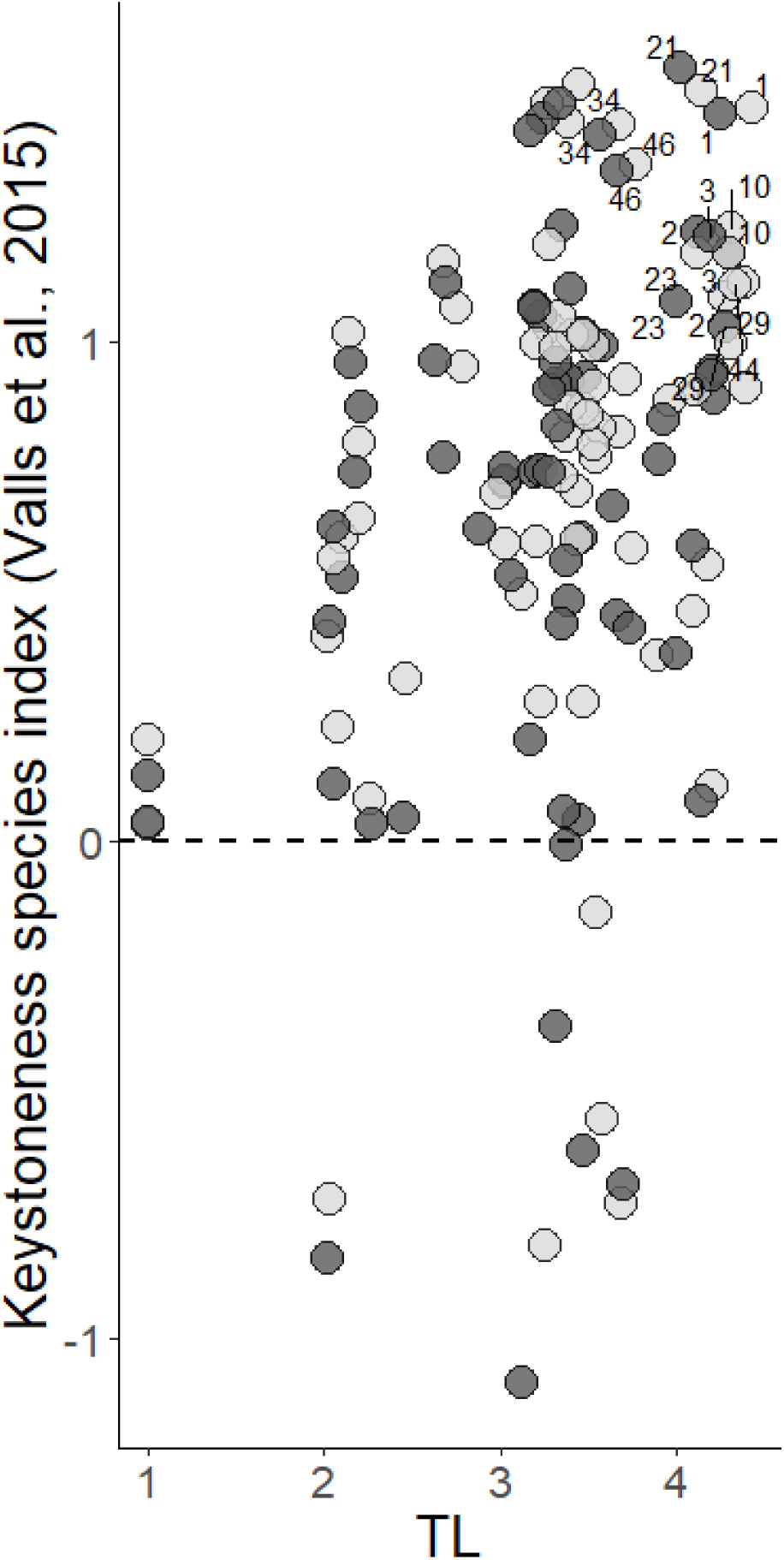
Keystone Index analysis of the continental slope of the Eastern Gulf of Lion Fisheries Restricted Area before its establishment (year 2008, light grey) and after (year 2016, dark grey). The numbers identify the functional group of the model (listed in Table A.1) with higher keystoneness index and relative total impact and trophic level (TL).

**Figure 5.**
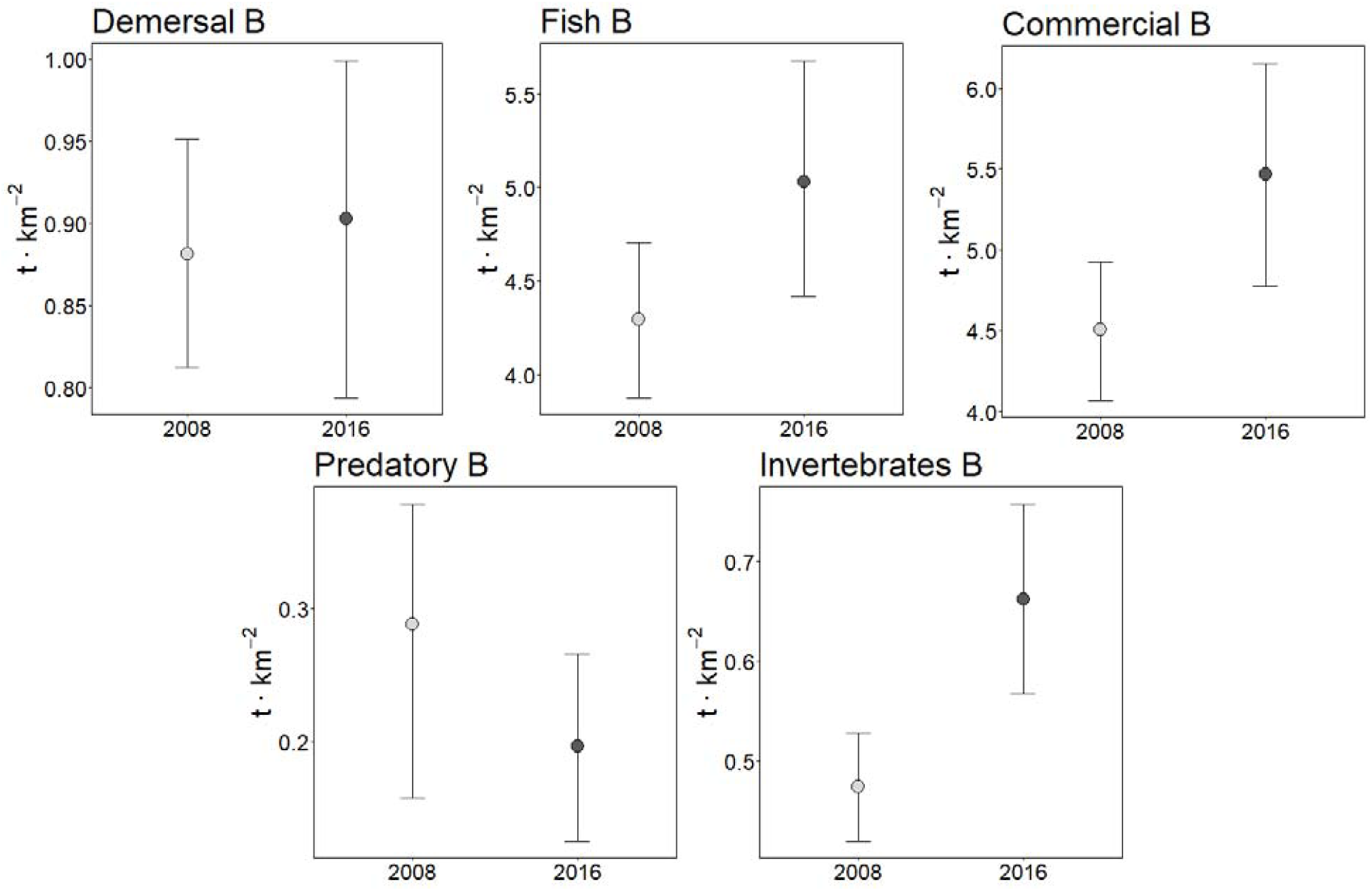
Biomass-based indicators of the continental slope of the Eastern Gulf of Lion Fisheries Restricted Area model before its establishment (year 2008, light grey) and after (year 2016, dark grey). B: Biomass. Error bars represent 95% confident intervals.

**Figure 6.**
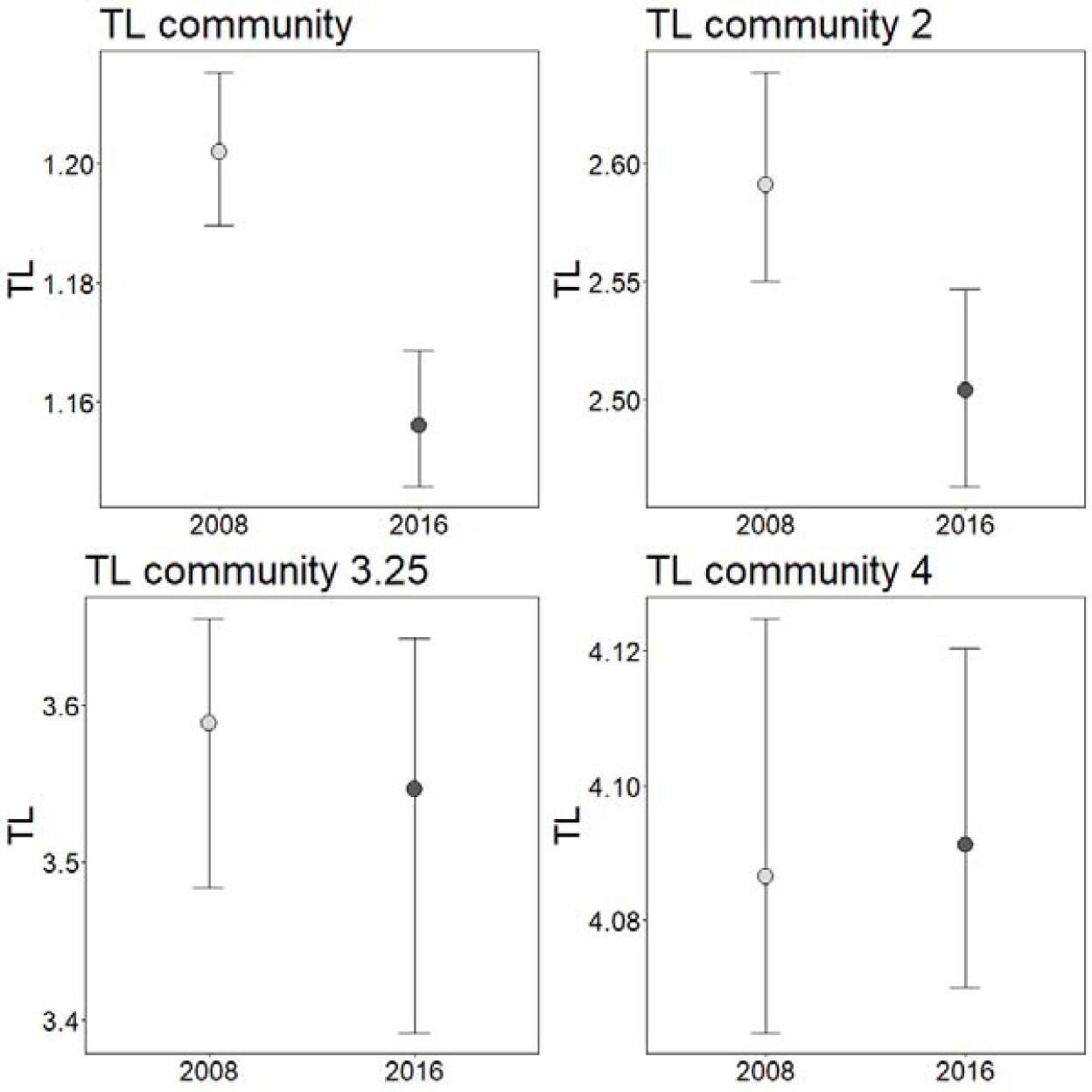
Trophic-based indicators of the continental slope of the Eastern Gulf of Lion Fisheries Restricted Area model before its establishment (year 2008, light grey) and after (year 2016, dark grey). TL: Trophic Level. Error bars represent 95% confident intervals.

**Figure 7.**
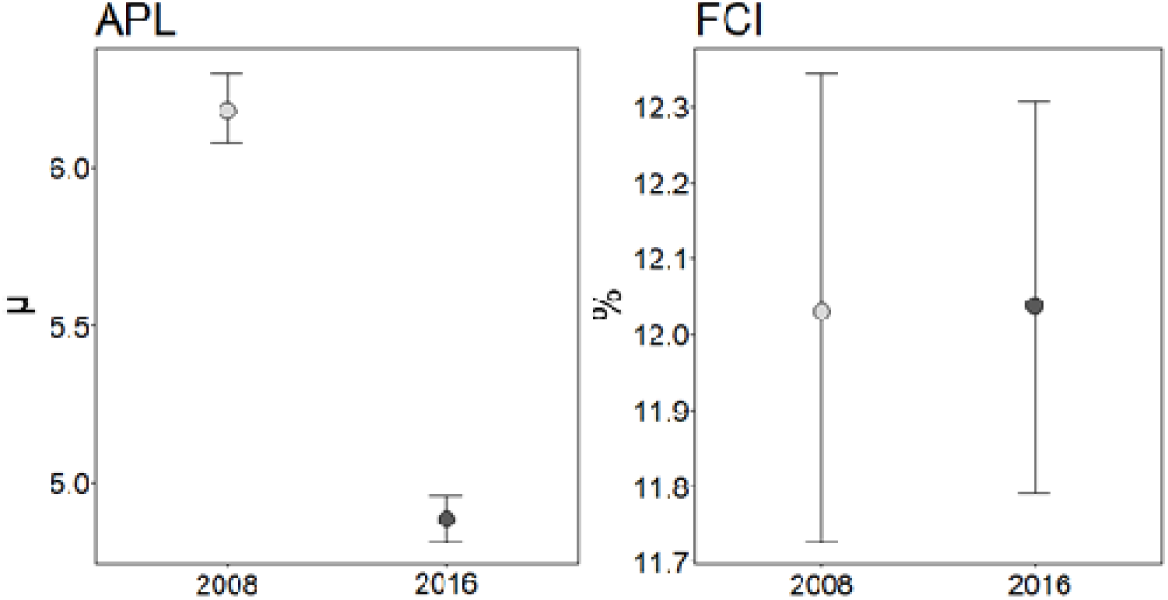
Flow-based indicators of the continental slope of the Eastern Gulf of Lion Fisheries Restricted Area model before its establishment (year 2008, light grey) and after (year 2016, dark grey). APL: The Average Path Length; FCI: The Finn’s Cycling Index. Error bars represent 95% confident intervals.

### 3.3. The impact of industrial fisheries

Total catch, Fish catch, and Discards increased once the FRA was establishment, while the TL of the catch and Demersal catch decreased (Figure 8). The MTI analysis applied to the industrial fisheries showed different patterns among fleets and CoSEGoL FRA states (2008 and 2016) (Figure 9). The highest positive impacting values were mostly found for the mid-water trawlers (MWT) (e.g. FG 12 (other large pelagic fish) or FG 55, (Norway lobster)). The most negative impacting values did not show any pattern among fleets. For example, while longliners (LON) impacted negatively on FG 11, 29 and 43 (swordfish, common dentex, and rays and skates, respectively), the bottom trawlers (BTW) impacted on FG 40, 51 and 52 (small-spotted catshark, Deep-water rose shrimp and blue and red shrimp, respectively) and the MTW on FG 10 (bluefin tuna). Although both models highlighted the same impacted FGs, the impact value of industrial fisheries over most FGs was different between both ecosystem states. Several FGs obtained lower positive values or higher negative values after the establishment of the CoSEGoL FRA. For example, other large pelagic fish (FG12) obtained an impacting value of 0.75 by MTW in 2008 while it was reduced to 0.64 in 2016, and blue and red shrimp (FG52) obtained an impacting value of −0.74 by BTW in 2008 while it increased to −0.81 in 2016.

**Figure 8.**
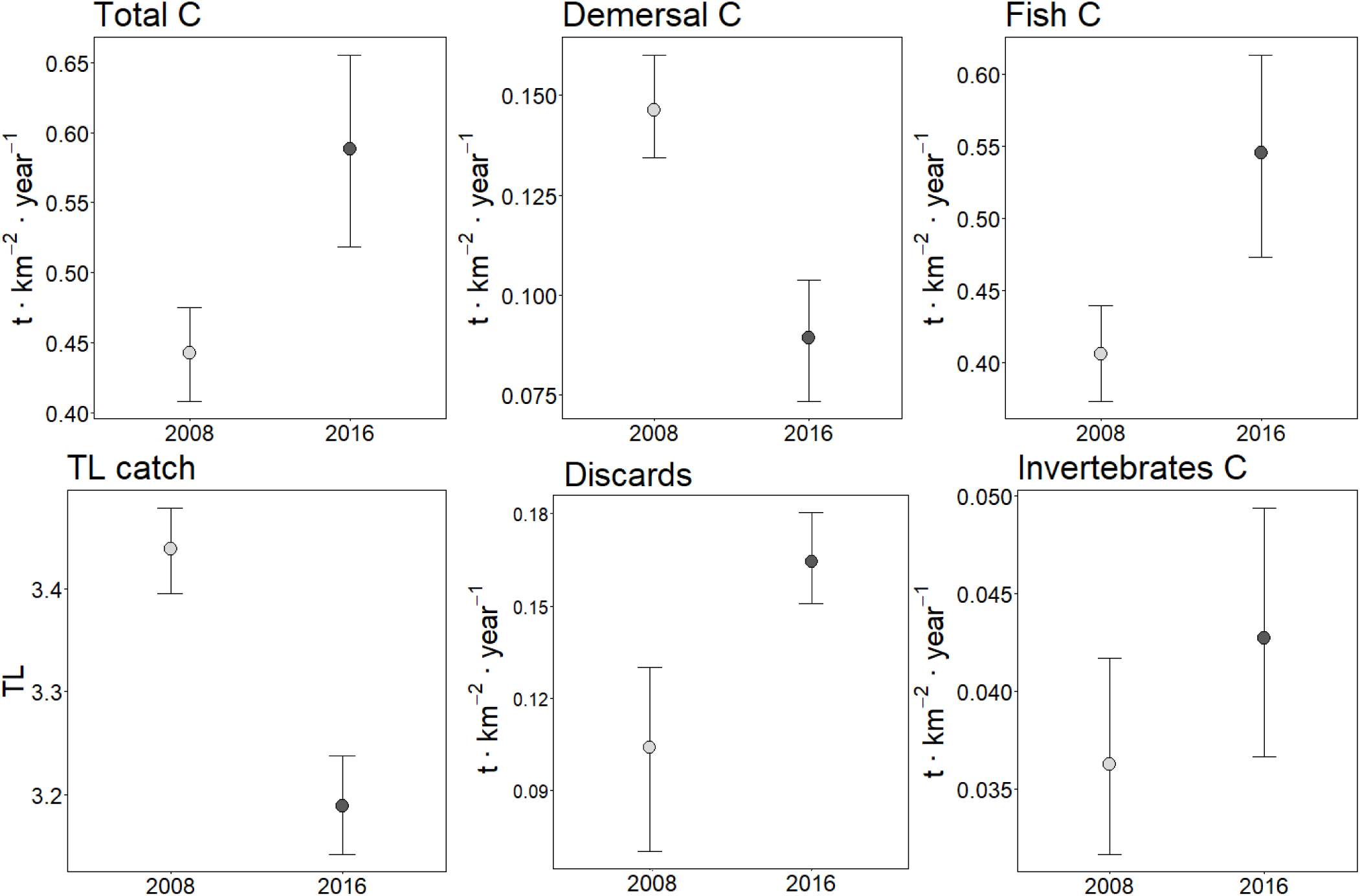
Catch-based indicators of the continental slope of the Eastern Gulf of Lion Fisheries Restricted Area model before its establishment (year 2008, light grey) and after (year 2016, dark grey). TL: Trophic Level, C: Catches. Error bars represent 95% confident intervals.

**Figure 9.**
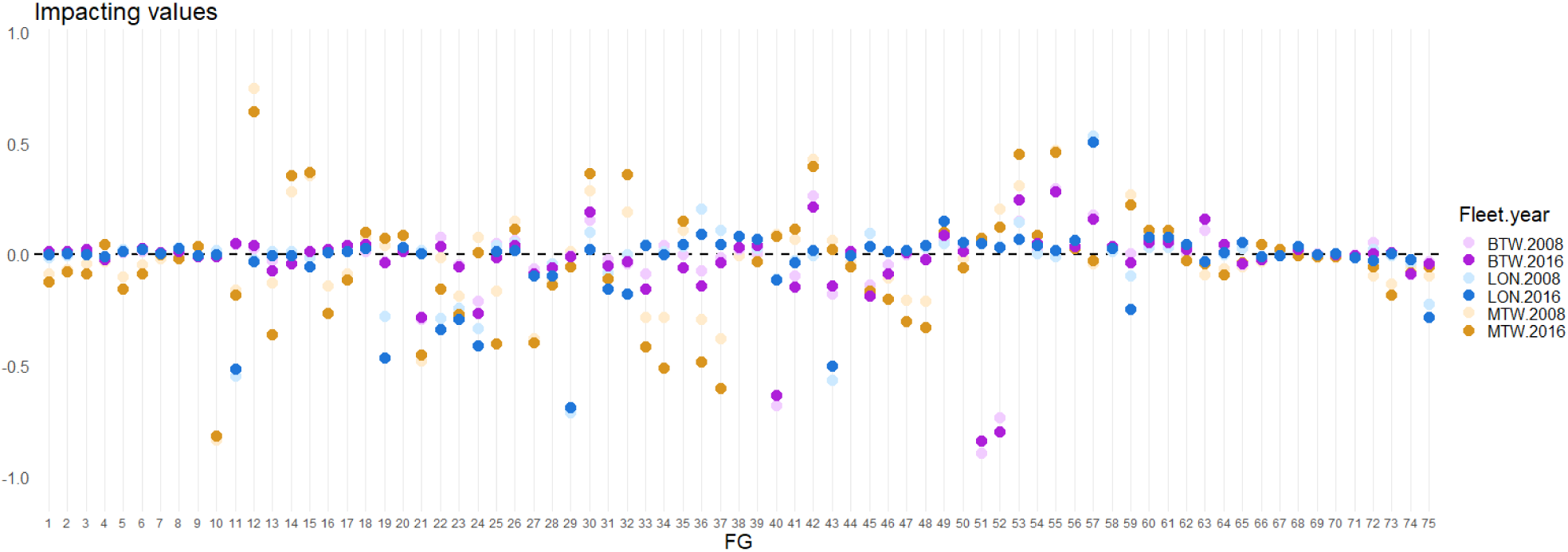
Industrial fisheries impacting values for each functional group of both ecosystem states of the continental slope of the Eastern Gulf of Lion Fisheries Restricted Area (2008 and 2016). X-axis identifies the FG number (except for 73, Midwater trawling: 74, Bottom trawling; and 75 Longliner) (MTW – Midwater trawling from France; BTW – bottom trawling from Spain and France; LON – Longliners from Spain).

### 3.4. Future scenarios of alternative management

Under baseline scenarios considering both RCP projections (BAU RCP4.5 and BAU RCP 8.5), the model predicted that European hake would decrease in both scenarios except in 2025 for scenario BAU RCP 4.5, while blue and red shrimp and Norway lobster showed decreasing biomass for both scenarios except in 2025 for scenario BAU RCP 8.5 (Figure 10). On the contrary, results showed an increase of biomass of anglerfish after 10 and 25 years of simulation (2025 and 2040, respectively) and blue whiting after 10 years (Figure 10). Within these scenarios, although biomass indicators increased in 2025, invertebrates, fish and commercial biomass indicators decreased in 2040 (Figure 11). Regarding catch-based indicators, invertebrates and demersal catch, and total catch and discards under RCP8.5 increased, while most indicators decreased in 2040 (Figure 12).

**Figure 10.**
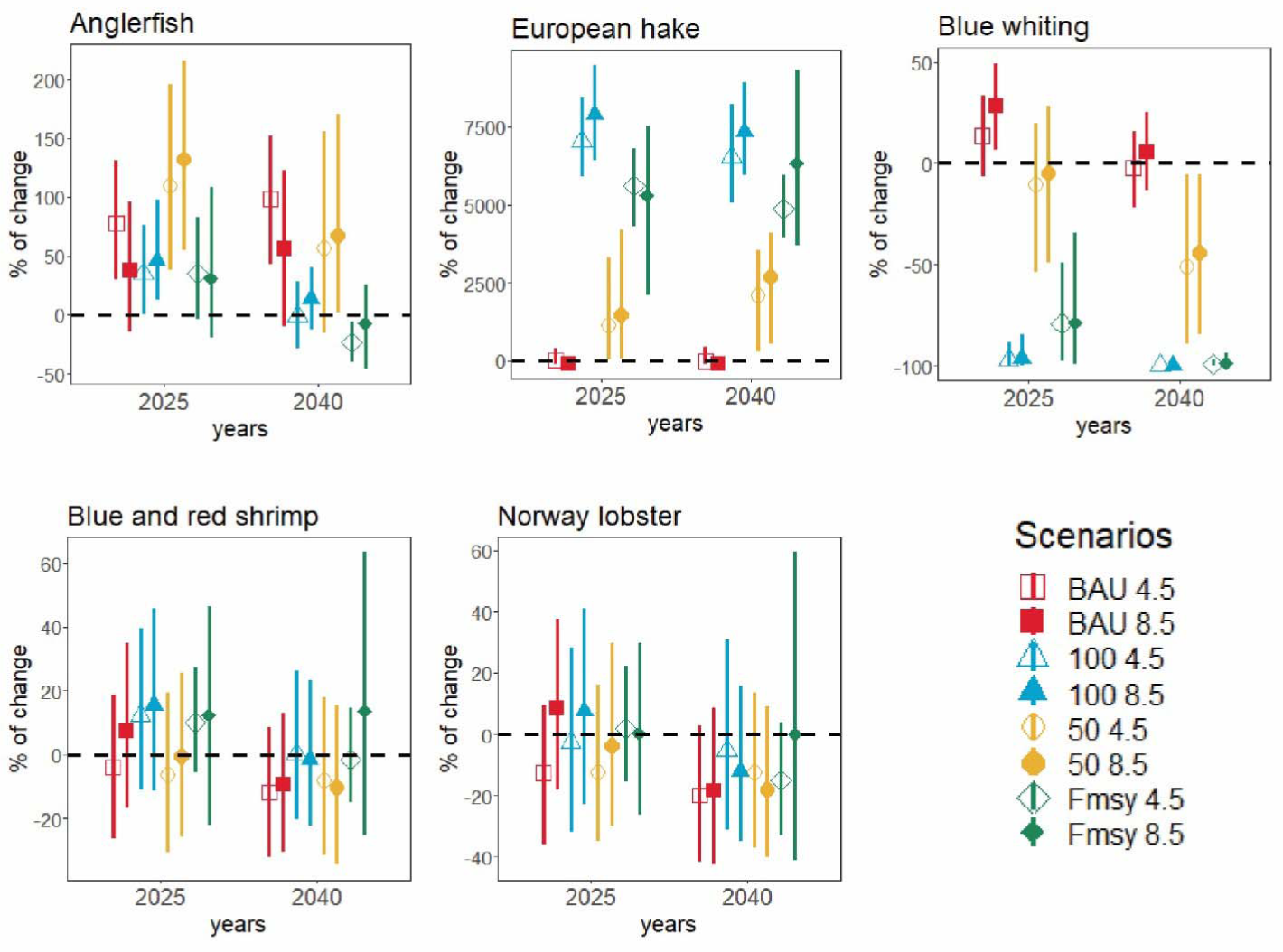
Mean percentage of change in biomass for targeted species under eight future scenarios of management of the continental slope of the Eastern Gulf of Lion Fisheries Restricted Area in 2025 and 2040. Error bars represent 95% confident intervals. Hollow points indicate scenarios under RCP4.5 projection, and solid points indicate scenarios under RCP8.5 projection.

**Figure 11.**
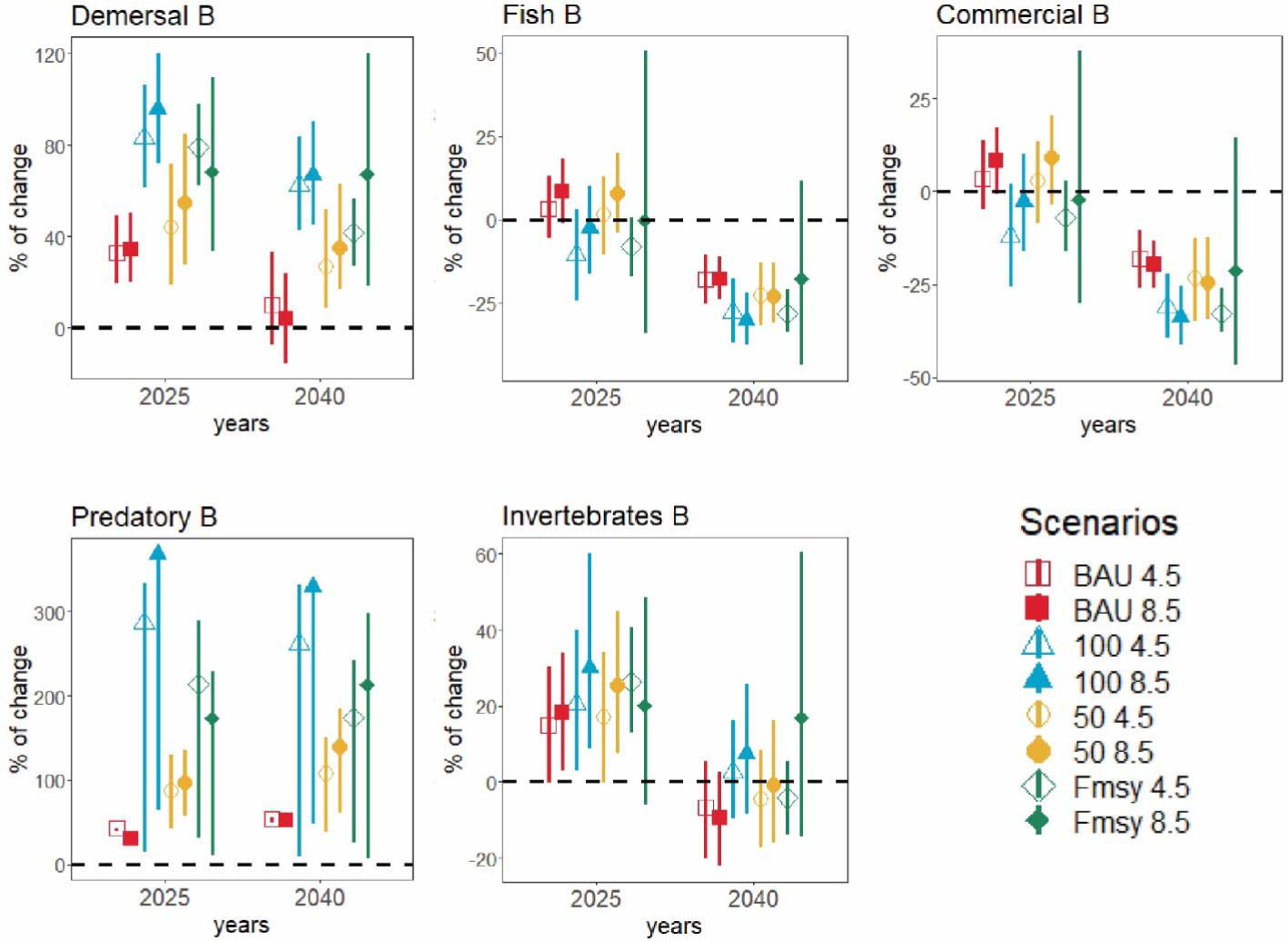
Mean percentage of change in biomass-based indicators under eight future scenarios of management of the continental slope of the Eastern Gulf of Lion Fisheries Restricted Area in 2025 and 2040. B: Biomass. Error bars represent 95% confident intervals. Hollow points indicate scenarios under RCP4.5 projection, and solid points indicate scenarios under RCP8.5 projection.

**Figure 12.**
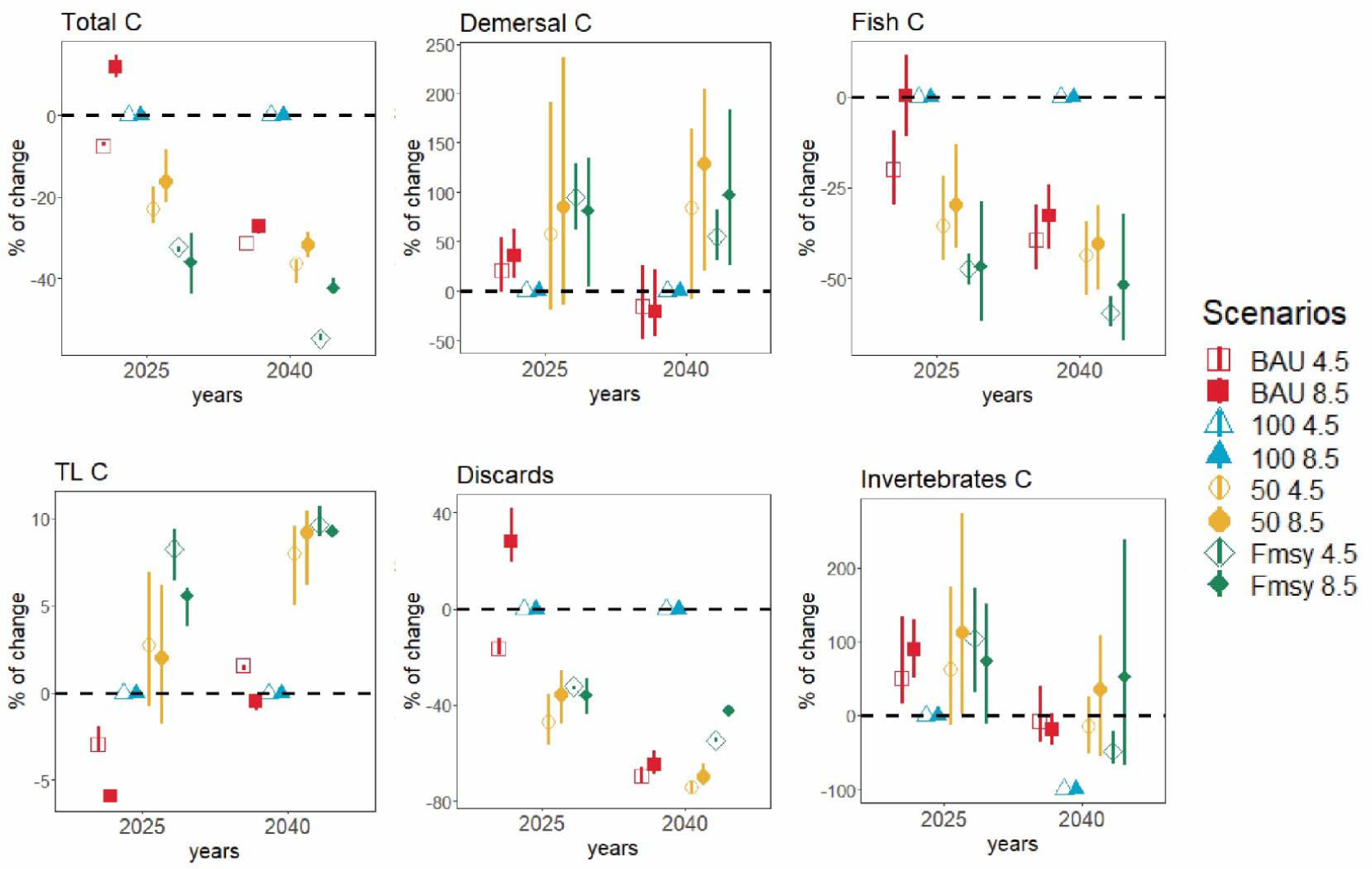
Mean percentage of change in catch-based indicators under eight future scenarios of management of the continental slope of the Eastern Gulf of Lion Fisheries Restricted Area in 2025 and 2040. TL: Trophic Level, C: Catches. Error bars represent 95% confident intervals. Hollow points indicate scenarios under RCP4.5 projection, and solid points indicate scenarios under RCP8.5 projection.

Applying a reduction of 100% on the fishing effort, both models with different RCP projections (scenarios 100 RCP 4.5 and 100 RCP 8.5) predicted increases on European hake biomass, and on anglerfish except for scenario 100 RCP 4.5 in 2040 (Figure 10), but predicted lower anglerfish biomass compared to scenarios BAU (Figure 10). On the contrary, blue whiting decreased as it did Norway lobster that decreased for both scenarios except for 100 RCP 8.5 in 2025. Blue and red shrimp increased on biomass for both scenarios except for 100 RCP 8.5 in 2040. We note that blue and red shrimp and Norway lobster obtained higher biomass predictions compared to baseline results except for Norway lobster for scenario 100 RCP 8.5 in 2025 (Figure 10).

Simulating a reduction of 50% of the fishing effort, the model predicted increases in the biomass of European hake, too. The percentage of change in biomass under scenarios 50 were higher than BAU scenarios but lower than scenarios 100. Results showed higher increase on anglerfish biomass than scenarios 100 and BAU, except in 2040 under RCP4.5 (Figure 10). In contrast, scenarios 50 RCP 4.5 and 50 RCP 8.5 predicted a decrease in biomass trends for blue whiting as scenario 100 but this reduction was smaller. Scenarios 50 RCP 4.5 and 50 RCP 8.5 also predicted a decreasing biomass trend for blue and red shrimp and Norway lobster. This pattern was similar to their baseline scenario predictions except for BAU RCP 8.5 in 2025 (Figure 10).

Under fishing at Maximum Sustainable Yield scenarios, both models (F_msy_ RCP 4.5 and F_msy_ RCP 8.5) predicted an increase in biomass trends for European hake, which was higher than baseline predictions. Scenarios F_msy_ RCP 4.5 and F_msy_ RCP 8.5 predicted an increase in 2025 and a decrease in 2040 for anglerfish biomass, respectively (Figure 10). These anglerfish predictions were lower than baseline predictions. Oppositely, scenario F_msy_ predicted a decreasing biomass trend for blue whiting, which was much lower than baseline projections (Figure 10). For blue and red shrimp and Norway lobster, these two models predicted an increase in biomass trends except for F_msy_ RCP 4.5 in 2040. Blue and red shrimp and Norway lobster predictions under scenario F_msy_ were higher than the baseline ones except for Norway lobster under RCP 8.5 in 2025 (Figure 10).

Under fishing scenarios, biomass-based indicators increased in 2025 except for the commercial and fish biomass indicators which decreased for scenario 100 and F_msy_ (Figure 11). In 2040, these indicators decreased except for demersal, predatory and invertebrates’ biomass. Scenarios 100 obtained higher biomass values than scenarios 50, except for fish and commercial biomass. Generally, most biomass-based scenarios showed higher mean values under RCP 8.5. Catch-based indicators showed decreasing trends for total and fish catch and discards, while invertebrates catch, demersal catch and trophic level of the catch increased (Figure 12). Most catch-based indicators obtained higher mean values under RCP8.5.

## 4. Discussion

Too often, fishing restricted areas are established without time-bound impact assessments and recovery indicators of success. Here, we presented a first attempt, to our knowledge, to develop a time dynamic ecosystem model of a fisheries restricted area in the Mediterranean Sea accounting for the effect of its establishment.

Overall, according to our results the CoSEGoL FRA failed at improving the condition of the ecosystem over time. Most ecological indicators showed higher values prior to the establishment of the FRA compared to after. The biomass-based indicators did not show positive effects of the establishment of the FRA on biomass of commercial, fish, predatory neither demersal community after eight years of protection in the study area. Trophic-based indicators showed a reduction in TL community and TL community 2 from 2008 to 2016, which could evidence an ecosystem degradation with time [4]. After the implementation of the CoSEGoL FRA, the APL decreased which could suggest higher stress, less maturity, and lower resilience of the ecosystem [57]. These results could show that the measure to freeze the fishing effort to 2008 levels established by GFCM was insufficient to allow the rebuilding and protection of demersal commercial stocks. This difference could also be due to a failure on the enforcement of the FRA and the consecutive degradation of the system over time due to higher impacts of fishing. In accordance to our results, a recent report developed in European waters [68] pointed out that the fleet operating in the Gulf of Lion is the one with the highest non-compliance rate regarding relative fishing power of the vessels. In addition, the Automatic Identification System (AIS) data provided by Global Fishing Watch [69] were recently used to demonstrate the illegal fishing activities inside several Mediterranean FRAs including the CoSEGoL FRA [70], documenting the lack of enforcement in these areas.

Our study showed that most FGs were highly impacted by industrial fisheries after the implementation of the FRA, and this impact was higher compared to the pre-establishment of the FRA. This pattern reinforces previous results and it is likely highlighting an increase in the impact of fisheries after the establishment of the FRA [12]. In accordance with this, most catch-based indicators increased in their values. The low effectiveness of the FRA was also identified through the fitting procedure of the ecosystem model to historical time series of data, which showed that the best model configuration was achieved when an annual increase of 10% on the fishing effort was included in the initial parameterization.

Our study also illustrates that future management simulations are useful to explore trade-offs on species’ recovery as well as potential effects at the ecosystem level. In general, baseline scenarios showed different biomass historical trends for target species, such as an increase in anglerfish and blue whiting with a decrease in European hake, blue and red shrimp and Norway lobster. These contrasting biomass trends suggested direct and indirect impacts of fisheries on the food-web, as seen in the MTI analysis. For instance, European hake is targeted by all fleets operating in the CoSEGoL FRA, especially by longliners, and a high negative impact is expected. Other negative biomass trends can be explained by the profound impacts of just one fleet, such as Norway lobster by bottom trawlers. Increasing biomass trends are due to multiple trophic effects triggered by decreases of various predators and competitors. Since baseline historical scenarios showed a lack of positive biomass trends for key target species (including European hake), these results are likely suggesting that fishing regulations established in 2008 have not been effective, in accordance with previous results and reports [68,70] and that management was not enough to achieve the CoSEGoL FRA objectives [14].

Alternative fishing management scenarios showed different biomass trends for target species. In general, none of these scenarios showed simultaneous biomass increases for all five target species. Even scenario 100, where all fishing activities were banned, failed to show recovery effects for all target species. This suggests an important role of trophic interactions between some of the targeted species in the demersal community. For instance, blue whiting is an important source of food for European hake [71] and as such when hake recovers it has a negative effect on its prey. Food-web models can represent a useful tool for MPA assessment that can help to identify ecological trade-offs and synergies [72]. Results also show that trade-offs must be considered between fisheries management and climate change [73] and emphasize the need to include other stressors than fisheries to appropriately assess the future of marine ecosystems [26,27].

Despite these trade-offs, overall, demersal and invertebrates’ biomass showed increasing trends with recovery scenarios, which indicates that the improvement on the status of demersal community may be possible under alternative management. In accordance with biomass-based indicators, catch-based indicators showed positive values for demersal and invertebrates catches. Additionally, although total catch and discards decreased, more substantial decreases in discards may indicate a move towards sustainable fishing under alternative future management scenarios. Our results also showed that the establishment of specific objectives should be the main aspect of implementing a restricted area to fisheries [74,75] and managers should focus on indicators related to the overall objective of this protection [76]. The CoSEGoL FRA was focused on demersal species [37], and our study showed that reducing fishing effort in the CoSEGoL FRA could benefit demersal species, in accordance to findings by other studies [18]. Regarding target species, biomass and catch-based indicators changed under different RCP scenarios in 2040, and thus climate change predictions under multiple scenarios should be considered for management purposes in the future [77].

During the study we dealt with several limitations. One was the lack of spatial-temporal series of catches and fleet distribution data, which could improve the analysis on the effect of the FRA potential benefits on the industrial fisheries. Although biomass data came from MEDITS survey database and are characterized spatially and temporally, catches within the FRA were assumed to be proportional to catches in the Geographical Sub-Area 7 (Gulf of Lion), scaled to fishing vessel presence at fishable velocities in the FRA. These assumptions increased the uncertainty in catch estimates. Additionally, the organization in charge to regulate the CoSEGoL FRA, the GFCM, reported the list of vessels operating in this area^2^, which differs from AIS data available at Global Fishing Watch [70]. Considering that enforcement has demonstrated to be an important feature to achieve ecological benefits in an MPA [78], this calls for a better understanding of the catch data inside the FRA and probably an improvement in the surveillance. Collecting time series of fishing activities inside the CoSEGoL FRA should be a monitoring priority, as previously highlighted for other marine protected areas [31,32,79]. In addition, response functions to sea temperature were included from a global database [64] because we lacked specific response functions in the study area. Specific sea temperature response function could improve the predictions under different RCP projections (e.g. [80]).

To our knowledge, this study represents the first attempt to develop a food-web model in an FRA and provides an assessment on the current management and potential outcomes of alternative fishing management scenarios. Our results suggest a failure on the recovery of target species in the restricted area under current management scenario. Results on future scenarios highlight the need to undertake important reductions in fishing effort in the FRA area, with highest benefits for marine resources and the ecosystem if the area would be closed to fishing. The CoSEGoL FRA could act as an important refugee of large spawners of commercial species which can contribute to rebuild demersal stocks in the Northwestern Mediterranean Sea [37]. However, the lack of enforcement and/or effectiveness of the FRA is contributing to its failure. The study also highlights the importance of considering trophic interactions when assessing the impacts of fishing management options, especially when target species are trophically related and include both predators and prey.

## Acknowledgments

This work was funded by EU Research Project SAFENET project (“Sustainable Fisheries in EU Mediterranean Waters through Network of MPAs.” Call for proposals MARE/2014/41, Grant Agreement n. 721708).

Additional Supplementary material may be found in the online version of this article:

**Appendix A.** Supplementary information about the ecosystem modelling approach.

**Appendix B.** Supplementary table: The CoSEGoL FRA FGs species composition and methods and references used to estimate the basic input parameters (Table B.1).

**Appendix C.** Supplementary table: Input parameters and outputs estimate for the CoSEGoL FRA model (Table C.1). Diet composition matrix for the CoSEGoL FRA model (Table C.2).

**Appendix D.** Supplementary tables: Confidence intervals used to describe the uncertainty for functional group (FG) and each input parameter of the balanced Ecopath model (Table D.1). Reference points used to develop the F_msy_ simulations for CoSEGoL FRA (Table D.2).

**Appendix E.** Supplementary figures: Historic and future trends under the two scenarios of IPCC projections of environmental variables considered in the CoSEGoL FRA model: sea water temperature (Figure E.1) and primary production (Figure E.2)

**Appendix F.** Supplementary table: Results of the fitting procedure of the CoSEGoL FRA ecosystem fitted to time series of data from 2008 to 2016 (Table F.1)

## Footnotes

1 http://www.criobe.pf/recherche/safenet/

2 *https://gfcmsitestorage.blob.core.windows.net/contents/DB/GoL/Html* and *http://www.fao.org/gfcm/data/fleet/fras*)

