## Supplemental Appendix B for "Current and potential contributions of the Gulf of Lion Fisheries Restricted Area to fisheries sustainability in the NW Mediterranean Sea"

Table B1. The continental slope of the Eastern Gulf of Lions (CoSEGoL) FRA functional groups (FGs) species composition and methods and references used to estimate the basic input parameters: Biomass (B) (t·km^-^²), Production over Biomass (year^-1^) (P/B), Consumption over Biomass (year^-1^) (Q/B), Diet (D), Catch (t·km^-^²·year^-1^) (C), Discard (t·km^-^²·year^-1^) (Di) and Ecotrophic Efficiency (year^-1^) (EE) of the Ecopath model in 2008.

| **Basic Input parameters** | **Method** | **Source** |
| --- | --- | --- |
| **1. Bottlenose dolphins (BOD):** *Tursiops truncatus* | | |
| **B** |  | [1–15] |
| **P/B** | Life history table | [16] |
| **Q/B** | From modified energy requirement equation: E =aW^0.714^ | [17–21] |
| **D** |  | [20] |
| **2. Striped dolphins (STD):** *Stenella coeruleoalba* | | |
| **B** |  | [3,6,6,9,22–32] |
| **P/B** | Life history table | [16] |
| **Q/B** | From modified energy requirement equation: E =aW^0.714^ | [17,18,21,33,34] |
| **D** |  | [35] |
| **3. Short-beaked common dolphin (COD):** *Delphinus delphis* | | |
| **B** |  | [6,26,36] |
| **P/B** | Life history table | [16] |
| **Q/B** | From modified energy requirement equation: E =aW^0.714^ | [17,18,21,37]^,^ |
| **D** |  | [38] |
| **4.** **Fin whale (FIW):** *Balaenoptera physalus* | | |
| **B** |  | [6,22,22,24,25,28,30,31,39,40] |
| **P/B** | Life history table | [16] |
| **Q/B** | From modified energy requirement equation: E =aW^0.714^ | [17,18,21] |
| **D** |  | [41] |
| **5.** **Deep sea-cetacean feeder marine mammals (DSM):** *Globicephala melas, Grampus griseus, Physeter macrocephalus, Ziphius cavirostris* | | |
| **B** |  | [3,6,9,22,42–51] |
| **P/B** | Life history table | [16] |
| **Q/B** | From modified energy requirement equation: E =aW^0.714^ | [17,18,21] |
| **D** |  | [22,42,52] |
| **6.** **Endangered and pelagic seabirds (ENS):** *Calonectris diomedea, Hydrobates pelagicus melitensis, Puffinus yelkouan, Puffinus mauretanicus* | | |
| **B** |  | [53–65] |
| **P/B** | Mortality (z)= -ln(Survival rate) | [66] |
| **Q/B** | Yearly food intake (including breeding and no breeding seasons)/B | [67,68] |
| **D** |  | [69–71] |
| **7.** **Loggerhead turtle (LGT):** *Caretta caretta* | | |
| **B** |  | [72–81] |
| **P/B** | Mortality (z)= -ln(Survival rate) | [66,81,82] |
| **Q/B** | Yearly food intake/B | [83] |
| **D** |  | [84] |
| **8.** **Pelagic sharks (PSK):** *Alopias vulpinus, Carcharhinus plumbeus, Carcharia taurus, Carcharodon carcharias, Galeorhinus galeus, Isurus oxyrinchus, Lamna nasus, Prionace glauca* | | |
| **B** |  | [75,85,86] and International Bottom Trawl Survey in the Mediterranean (MEDITS) [87] |
| **P/B** | $\log M=-0.0066-0.279\cdot\log L_{\infty}+0.6543\cdot\log k+0.4634\cdot\log T$  Mortality (z)= F+M | Empirical equation from [88]; L_∞_ and k parameter from [89–92] |
| **Q/B** | $\log Q/B=6.37-1.5045\cdot\log T^{'}-0.168\cdot\log W_{\infty}+0.1399\cdot Pf+0.2765\cdot HD$ | Empirical equation from [93]; *a* and *b* parameters from [94–97] |
| **D** |  | [98] |
| **9.** **Non-commercial large pelagic fish (NLP):** *Cetorhinus maximus, Mobula mobular, Mola mola* | | |
| **B** |  | [75,85,86] and International Bottom Trawl Survey in the Mediterranean (MEDITS) [87] |
| **P/B** | $\log M=-0.0066-0.279\cdot\log L_{\infty}+0.6543\cdot\log k+0.4634\cdot\log T$  Mortality (z)= F+M | Empirical equation from [88]; L_∞_ and k parameter from [99–101] |
| **Q/B** | $\log Q/B=6.37-1.5045\cdot\log T^{'}-0.168\cdot\log W_{\infty}+0.1399\cdot Pf+0.2765\cdot HD$ | Empirical equation from [93]; *a* and *b* parameters from [102–104] |
| **D** |  | [105] |
| **10.** **Bluefin tuna (BLF):** *Thunnus thynnus* | | |
| **B** |  | [106] |
| **P/B** | $\log M=-0.0066-0.279\cdot\log L_{\infty}+0.6543\cdot\log k+0.4634\cdot\log T$  Mortality (z)= F+M | Empirical equation from [88]; L_∞_ and k parameter from [107] |
| **Q/B** | $\log Q/B=6.37-1.5045\cdot\log T^{'}-0.168\cdot\log W_{\infty}+0.1399\cdot Pf+0.2765\cdot HD$ | Empirical equation from [93]; *a* and *b* parameters from [108] |
| **D** |  | [109] |
| **B-C** |  | [110,111] |
| 11. **Swordfish (SWO):** *Xiphias gladius* | | |
| **B** |  | [112] |
| **P/B** | $\log M=-0.0066-0.279\cdot\log L_{\infty}+0.6543\cdot\log k+0.4634\cdot\log T$  Mortality (z)= F+M | Empirical equation from [88]; L_∞_ and k parameter from [113] |
| **Q/B** | $\log Q/B=6.37-1.5045\cdot\log T^{'}-0.168\cdot\log W_{\infty}+0.1399\cdot Pf+0.2765\cdot HD$ | Empirical equation from [93]; *a* and *b* parameters from [114] |
| **D** |  | [84] |
| **B-C** |  | [110,111] |
| **12.** **Other large pelagic fish (OLP):** *Coryphaena hippurus, Lichia amia, Seriola dumerili,* | | |
| **B** |  | [115] and International Bottom Trawl Survey in the Mediterranean (MEDITS) [87] |
| **P/B** | $\log M=-0.0066-0.279\cdot\log L_{\infty}+0.6543\cdot\log k+0.4634\cdot\log T$  Mortality (z)= F+M | Empirical equation from [88]; L_∞_ and k parameter from [116,117] |
| **Q/B** | $\log Q/B=6.37-1.5045\cdot\log T^{'}-0.168\cdot\log W_{\infty}+0.1399\cdot Pf+0.2765\cdot HD$ | Empirical equation from [93]; *a* and *b* parameters from [116,117] |
| **D** |  | [118–120] |
| **C** |  | [110,111] |
| **13.** **Mackerels (MCK):** *Scomber scombrus, Scomber colias* | | |
| **B** |  | International Bottom Trawl Survey in the Mediterranean (MEDITS) |
| **P/B** | $\log M=-0.0066-0.279\cdot\log L_{\infty}+0.6543\cdot\log k+0.4634\cdot\log T$  Mortality (z)= F+M | Empirical equation from [88]; L_∞_ and k parameter from [121,122] |
| **Q/B** | $\log Q/B=6.37-1.5045\cdot\log T^{'}-0.168\cdot\log W_{\infty}+0.1399\cdot Pf+0.2765\cdot HD$ | Empirical equation from [93]; *a* and *b* parameters from [123,124] |
| **D** |  | [84,125] |
| **C** |  | [110,111] |
| **14.** **Horse mackerels (HRM):** *Trachurus trachurus, Trachurus mediterraneus, Trachurus picturatus* | | |
| **B** |  | International Bottom Trawl Survey in the Mediterranean (MEDITS) [87] |
| **P/B** | $\log M=-0.0066-0.279\cdot\log L_{\infty}+0.6543\cdot\log k+0.4634\cdot\log T$  Mortality (z)= F+M | Empirical equation from [88]; L_∞_ and k parameter from [126,127] |
| **Q/B** | $\log Q/B=6.37-1.5045\cdot\log T^{'}-0.168\cdot\log W_{\infty}+0.1399\cdot Pf+0.2765\cdot HD$ | Empirical equation from [93]; *a* and *b* parameters from [123] |
| **D** |  | [84,128] |
| **C** |  | [110,111] |
| **15.** **Other medium pelagic fish (OMP):** *Alosa alosa, Alosa fallax, Caranx rhonchus, Pomatomus saltatrix, Sarda sarda, Scomberesox saurus, Sphyraena sphyraena* | | |
| **B** |  | International Bottom Trawl Survey in the Mediterranean (MEDITS) |
| **P/B** | $\log M=-0.0066-0.279\cdot\log L_{\infty}+0.6543\cdot\log k+0.4634\cdot\log T$  Mortality (z)= F+M | Empirical equation from [88]; L_∞_ and k parameter from [103,129–133] |
| **Q/B** | $\log Q/B=6.37-1.5045\cdot\log T^{'}-0.168\cdot\log W_{\infty}+0.1399\cdot Pf+0.2765\cdot HD$ | Empirical equation from [93]; *a* and *b* parameters from [103,114,124,132,134,135] |
| **D** |  | [136–140] |
| **C** |  | [110,111] |
| **16.** **European sardine (ESA):** *Sardina pilchardus* | | |
| **B** |  | Mediterranean International Acoustic Survey (MEDIAS) [141] |
| **P/B** | $\log M=-0.0066-0.279\cdot\log L_{\infty}+0.6543\cdot\log k+0.4634\cdot\log T$  Mortality (z)= F+M | Empirical equation from [88]; L_∞_ and k parameter from [142] |
| **Q/B** | $\log Q/B=6.37-1.5045\cdot\log T^{'}-0.168\cdot\log W_{\infty}+0.1399\cdot Pf+0.2765\cdot HD$ | Empirical equation from [93]; *a* and *b* parameters from  Joint Research Centre, DCF, Data Collection Framework (https://datacollection.jrc.ec.europa.eu/) |
| **D** |  | [143] |
| **C** |  | [110,111] |
| **17.** **European anchovy (EAN):** *Engraulis encrasicolus* | | |
| **B** |  | Mediterranean International Acoustic Survey (MEDIAS) [141] |
| **P/B** | $\log M=-0.0066-0.279\cdot\log L_{\infty}+0.6543\cdot\log k+0.4634\cdot\log T$  Mortality (z)= F+M | Empirical equation from [88]; L_∞_ and k parameter from [144] |
| **Q/B** | $\log Q/B=6.37-1.5045\cdot\log T^{'}-0.168\cdot\log W_{\infty}+0.1399\cdot Pf+0.2765\cdot HD$ | Empirical equation from [93]; *a* and *b* parameters from  Joint Research Centre, DCF, Data Collection Framework (https://datacollection.jrc.ec.europa.eu/) |
| **D** |  | [145] |
| **C** |  | [110,111] |
| **18.** **Other small pelagic fish (OSP):** *Spicara flexuosa, Spicara maena, Spicara smaris, Sprattus sprattus* | | |
| **B** |  | International Bottom Trawl Survey in the Mediterranean (MEDITS) [87] |
| **P/B** | $\log M=-0.0066-0.279\cdot\log L_{\infty}+0.6543\cdot\log k+0.4634\cdot\log T$  Mortality (z)= F+M | Empirical equation from [88]; L_∞_ and k parameter from [146–148] |
| **Q/B** | $\log Q/B=6.37-1.5045\cdot\log T^{'}-0.168\cdot\log W_{\infty}+0.1399\cdot Pf+0.2765\cdot HD$ | Empirical equation from [93]; *a* and *b* parameters from [123,149] |
| **D** |  | [128,150,151] |
| **C** |  | [110,111] |
| **19.** **Benthopelagic fish (BTP):** *Coelorinchus caelorhincus, Hymenocephalus italicus, Lepidion lepidion, Lepidopus caudatus, Nezumia sclerorhynchus* | | |
| **B** |  | International Bottom Trawl Survey in the Mediterranean (MEDITS) [87] |
| **P/B** | $\log M=-0.0066-0.279\cdot\log L_{\infty}+0.6543\cdot\log k+0.4634\cdot\log T$  Mortality (z)= F+M | Empirical equation from [88]; L_∞_ and k parameter from [117,152–154] |
| **Q/B** | $\log Q/B=6.37-1.5045\cdot\log T^{'}-0.168\cdot\log W_{\infty}+0.1399\cdot Pf+0.2765\cdot HD$ | Empirical equation from [93]; *a* and *b* parameters from [117,123,155,156] |
| **D** |  | [157–160] |
| **C** |  | [110,111] |
| **20.** **No commercial meso (bathy) pelagic fish (NBTP):** *Argyropelecus hemigymnus, Gadiculus argenteus* | | |
| **B** |  | International Bottom Trawl Survey in the Mediterranean (MEDITS) [87] |
| **P/B** | $\log M=-0.0066-0.279\cdot\log L_{\infty}+0.6543\cdot\log k+0.4634\cdot\log T$  Mortality (z)= F+M | Empirical equation from [88]; L_∞_ and k parameter from [117,161] |
| **Q/B** | $\log Q/B=6.37-1.5045\cdot\log T^{'}-0.168\cdot\log W_{\infty}+0.1399\cdot Pf+0.2765\cdot HD$ | Empirical equation from [93]; *a* and *b* parameters from [117,162] |
| **D** |  | [159,163] |
| **Di** |  | [110,111] |
| **21.** **Anglerfish (ANG):** *Lophius budegassa, Lophius piscatorius* | | |
| **B** |  | International Bottom Trawl Survey in the Mediterranean (MEDITS) [87] |
| **P/B** | $\log M=-0.0066-0.279\cdot\log L_{\infty}+0.6543\cdot\log k+0.4634\cdot\log T$  Mortality (z)= F+M | Empirical equation from [88]; L_∞_ and k parameter from [164,165] |
| **Q/B** | $\log Q/B=6.37-1.5045\cdot\log T^{'}-0.168\cdot\log W_{\infty}+0.1399\cdot Pf+0.2765\cdot HD$ | Empirical equation from [93]; *a* and *b* parameters from [123,125] |
| **D** |  | [159,166] |
| **C** |  | [110,111] |
| **22.** **European conger (ECO):** *Conger conger* | | |
| **B** |  | International Bottom Trawl Survey in the Mediterranean (MEDITS) [87] |
| **P/B** | $\log M=-0.0066-0.279\cdot\log L_{\infty}+0.6543\cdot\log k+0.4634\cdot\log T$  Mortality (z)= F+M | Empirical equation from [88]; L_∞_ and k parameter from [121] |
| **Q/B** | $\log Q/B=6.37-1.5045\cdot\log T^{'}-0.168\cdot\log W_{\infty}+0.1399\cdot Pf+0.2765\cdot HD$ | Empirical equation from [93]; *a* and *b* parameters from [123] |
| **D** |  | [159] |
| **C** |  | [110,111] |
| **23.** **European hake (EHK):** *Merluccius merluccius* | | |
| **B** |  | International Bottom Trawl Survey in the Mediterranean (MEDITS) [87] |
| **P/B** | $\log M=-0.0066-0.279\cdot\log L_{\infty}+0.6543\cdot\log k+0.4634\cdot\log T$  Mortality (z)= F+M | Empirical equation from [88]; L_∞_ and k parameter from [121] |
| **Q/B** | $\log Q/B=6.37-1.5045\cdot\log T^{'}-0.168\cdot\log W_{\infty}+0.1399\cdot Pf+0.2765\cdot HD$ | Empirical equation from [93]; *a* and *b* parameters from [167] |
| **D** |  | [168] |
| **C** |  | [110,111] |
| **24.** **Other commercial large demersal fish (OLD):** *Molva macrophthalma* | | |
| **B** |  | International Bottom Trawl Survey in the Mediterranean (MEDITS) [87] |
| **P/B** | $\log M=-0.0066-0.279\cdot\log L_{\infty}+0.6543\cdot\log k+0.4634\cdot\log T$  Mortality (z)= F+M | Empirical equation from [88]; L_∞_ and k parameter from [169] |
| **Q/B** | $\log Q/B=6.37-1.5045\cdot\log T^{'}-0.168\cdot\log W_{\infty}+0.1399\cdot Pf+0.2765\cdot HD$ | Empirical equation from [93]; *a* and *b* parameters from [123] |
| **D** |  | [159] |
| **C** |  | [110,111] |
| **25.** **Poor cod (PCO):** *Trisopterus capelanus* | | |
| **B** |  | International Bottom Trawl Survey in the Mediterranean (MEDITS) |
| **P/B** | $\log M=-0.0066-0.279\cdot\log L_{\infty}+0.6543\cdot\log k+0.4634\cdot\log T$  Mortality (z)= F+M | Empirical equation from [88]; L_∞_ and k parameter from [170] |
| **Q/B** | $\log Q/B=6.37-1.5045\cdot\log T^{'}-0.168\cdot\log W_{\infty}+0.1399\cdot Pf+0.2765\cdot HD$ | Empirical equation from [93]; *a* and *b* parameters from [123] |
| **D** |  | [171] |
| **C** |  | [110,111] |
| **26.** **Blue whiting (BLW):** *Micromesistius poutassou* | | |
| **B** |  | International Bottom Trawl Survey in the Mediterranean (MEDITS) [87] |
| **P/B** | $\log M=-0.0066-0.279\cdot\log L_{\infty}+0.6543\cdot\log k+0.4634\cdot\log T$  Mortality (z)= F+M | Empirical equation from [88]; L_∞_ and k parameter from [172] |
| **Q/B** | $\log Q/B=6.37-1.5045\cdot\log T^{'}-0.168\cdot\log W_{\infty}+0.1399\cdot Pf+0.2765\cdot HD$ | Empirical equation from [93]; *a* and *b* parameters from [173] |
| **D** |  | [159] |
| **C** |  | [110,111] |
| **27.** **Common pandora (CPA):** *Pagellus erythrinus* | | |
| **B** |  | International Bottom Trawl Survey in the Mediterranean (MEDITS) [87] |
| **P/B** | $\log M=-0.0066-0.279\cdot\log L_{\infty}+0.6543\cdot\log k+0.4634\cdot\log T$  Mortality (z)= F+M | Empirical equation from [88]; L_∞_ and k parameter from [174] |
| **Q/B** | $\log Q/B=6.37-1.5045\cdot\log T^{'}-0.168\cdot\log W_{\infty}+0.1399\cdot Pf+0.2765\cdot HD$ | Empirical equation from [93]; *a* and *b* parameters from [121] |
| **D** |  |  |
| **C** |  | [110,111] |
| **28.** **Sparidae (SPA):** *Boops boops, Diplodus annularis, Pagellus acarne, Pagellus bogaraveo,* | | |
| **B** |  | International Bottom Trawl Survey in the Mediterranean (MEDITS) [87] |
| **P/B** | $\log M=-0.0066-0.279\cdot\log L_{\infty}+0.6543\cdot\log k+0.4634\cdot\log T$  Mortality (z)= F+M | Empirical equation from [88]; L_∞_ and k parameter from [174–176] |
| **Q/B** | $\log Q/B=6.37-1.5045\cdot\log T^{'}-0.168\cdot\log W_{\infty}+0.1399\cdot Pf+0.2765\cdot HD$ | Empirical equation from [93]; *a* and *b* parameters from [121,123] |
| **D** |  | [128,136,177] |
| **C** |  | [110,111] |
| **29. Common dentex (DEN):** *Dentex dentex* | | |
| **B** |  | International Bottom Trawl Survey in the Mediterranean (MEDITS) [87] |
| **P/B** | $\log M=-0.0066-0.279\cdot\log L_{\infty}+0.6543\cdot\log k+0.4634\cdot\log T$  Mortality (z)= F+M | Empirical equation from [88]; L_∞_ and k parameter from [174] |
| **Q/B** | $\log Q/B=6.37-1.5045\cdot\log T^{'}-0.168\cdot\log W_{\infty}+0.1399\cdot Pf+0.2765\cdot HD$ | Empirical equation from [93]; *a* and *b* parameters from [178] |
| **D** |  | [178] |
| **C** |  | [110,111] |
| **30. Red scorpionfish (RSC):** *Scorpaena scrofa* | | |
| **B** |  | International Bottom Trawl Survey in the Mediterranean (MEDITS) [87] |
| **P/B** | $\log M=-0.0066-0.279\cdot\log L_{\infty}+0.6543\cdot\log k+0.4634\cdot\log T$  Mortality (z)= F+M | Empirical equation from [88]; L_∞_ and k parameter from [121,146,179–184] |
| **Q/B** | $\log Q/B=6.37-1.5045\cdot\log T^{'}-0.168\cdot\log W_{\infty}+0.1399\cdot Pf+0.2765\cdot HD$ | Empirical equation from [93]; *a* and *b* parameters from [117] |
| **D** |  | [128] |
| **C** |  | [110,111] |
| **31.** **Scorpaenidae (SCO):** *Chelidonichthys cuculus, Eutrigla gurnardus, Helicolenus dactylopterus, Lepidotrigla cavillone, Lepidotrigla dieuzeidei, Scorpaena elongata, Scorpaena notata, Trigla lyra, Trigloporus lastoviza* | | |
| **B** |  | International Bottom Trawl Survey in the Mediterranean (MEDITS) |
| **P/B** | $\log M=-0.0066-0.279\cdot\log L_{\infty}+0.6543\cdot\log k+0.4634\cdot\log T$  Mortality (z)= F+M | Empirical equation from [88]; L_∞_ and k parameter from [121,146,179–184] |
| **Q/B** | $\log Q/B=6.37-1.5045\cdot\log T^{'}-0.168\cdot\log W_{\infty}+0.1399\cdot Pf+0.2765\cdot HD$ | Empirical equation from [93]; *a* and *b* parameters from [102,121,123] |
| **D** |  | [128,159,181,185–189] |
| **C** |  | [110,111] |
| **32.** **Labridae and serranidae (LAS):** *Serranus cabrilla, Serranus hepatus* | | |
| **B** |  | International Bottom Trawl Survey in the Mediterranean (MEDITS) [87] |
| **P/B** | $\log M=-0.0066-0.279\cdot\log L_{\infty}+0.6543\cdot\log k+0.4634\cdot\log T$  Mortality (z)= F+M | Empirical equation from [88]; L_∞_ and k parameter from [117,123] |
| **Q/B** | $\log Q/B=6.37-1.5045\cdot\log T^{'}-0.168\cdot\log W_{\infty}+0.1399\cdot Pf+0.2765\cdot HD$ | Empirical equation from [93]; *a* and *b* parameters from [123] |
| **D** |  | [128,185] |
| **C** |  | [110,111] |
| **33.** **Flatfish (FLA):** *Arnoglossus laterna, Arnoglossus rueppelii, Arnoglossus thori, Citharus linguatula, Lepidorhombus boscii, Lepidorhombus whiffiagonis, Microchirus variegatus, Solea solea, Symphurus nigrescens,* | | |
| **B** |  | International Bottom Trawl Survey in the Mediterranean (MEDITS) |
| **P/B** | $\log M=-0.0066-0.279\cdot\log L_{\infty}+0.6543\cdot\log k+0.4634\cdot\log T$  Mortality (z)= F+M | Empirical equation from [88]; *a* and *b* parameters from [117,190–193] |
| **Q/B** | $\log Q/B=6.37-1.5045\cdot\log T^{'}-0.168\cdot\log W_{\infty}+0.1399\cdot Pf+0.2765\cdot HD$ | Empirical equation from [93]; *a* and *b* parameters from [117,123,194] |
| **D** |  | [128,159,185,195,196] |
| **C** |  | [110,111] |
| **34.** **Other commercial medium demersal fish (CMD):** *Gaidropsarus biscayensis, Gaidropsarus mediterraneus, Phycis phycis, Zeus faber* | | |
| **B** |  | International Bottom Trawl Survey in the Mediterranean (MEDITS) [87] |
| **P/B** | $\log M=-0.0066-0.279\cdot\log L_{\infty}+0.6543\cdot\log k+0.4634\cdot\log T$  Mortality (z)= F+M | Empirical equation from [88]; L_∞_ and k parameter from [117,193,197] |
| **Q/B** | $\log Q/B=6.37-1.5045\cdot\log T^{'}-0.168\cdot\log W_{\infty}+0.1399\cdot Pf+0.2765\cdot HD$ | Empirical equation from [93]; *a* and *b* parameters from [121,123,198] |
| **D** |  | [159,185,199] |
| **C** |  | [110,111] |
| **35.** **No commercial medium demersal fish (NMD):** *Aulopus filamentosus,* *Cepola macrophthalma, Ophisurus serpens* | | |
| **B** |  | International Bottom Trawl Survey in the Mediterranean (MEDITS) [87] |
| **P/B** | $\log M=-0.0066-0.279\cdot\log L_{\infty}+0.6543\cdot\log k+0.4634\cdot\log T$  Mortality (z)= F+M | Empirical equation from [88]; L_∞_ and k parameter from [200,201] |
| **Q/B** | $\log Q/B=6.37-1.5045\cdot\log T^{'}-0.168\cdot\log W_{\infty}+0.1399\cdot Pf+0.2765\cdot HD$ | Empirical equation from [93]; *a* and *b* parameters from [200,202,203] |
| **D** |  | [136,204] |
| **Di** |  | [110,111] |
| 36. **Red mullet (RMU):** *Mullus barbatus* | | |
| **B** |  | International Bottom Trawl Survey in the Mediterranean (MEDITS) [87] |
| **P/B** | $\log M=-0.0066-0.279\cdot\log L_{\infty}+0.6543\cdot\log k+0.4634\cdot\log T$  Mortality (z)= F+M | Empirical equation from [88]; L_∞_ and k parameter from [205] |
| **Q/B** | $\log Q/B=6.37-1.5045\cdot\log T^{'}-0.168\cdot\log W_{\infty}+0.1399\cdot Pf+0.2765\cdot HD$ | Empirical equation from [93]; *a* and *b* parameters from  Joint Research Centre, DCF, Data Collection Framework (https://datacollection.jrc.ec.europa.eu/) |
| **D** |  | [206] |
| **C** |  | [110,111] |
| 37. **Surmullet (SUR):** *Mullus surmuletus* | | |
| **B** |  | International Bottom Trawl Survey in the Mediterranean (MEDITS) [87] |
| **P/B** | $\log M=-0.0066-0.279\cdot\log L_{\infty}+0.6543\cdot\log k+0.4634\cdot\log T$  Mortality (z)= F+M | Empirical equation from [88]; L_∞_ and k parameter from [121] |
| **Q/B** | $\log Q/B=6.37-1.5045\cdot\log T^{'}-0.168\cdot\log W_{\infty}+0.1399\cdot Pf+0.2765\cdot HD$ | Empirical equation from [93]; *a* and *b* parameters from  Joint Research Centre, DCF, Data Collection Framework (https://datacollection.jrc.ec.europa.eu/) |
| **D** |  | [128] |
| **C** |  | [110,111] |
| 38. **No** **commercial small demersal fish (NSD):** *Blennius ocellaris, Callionymus maculatus, Capros aper, Chlorophthalmus agassizi, Deltentosteus quadrimaculatus, Lesueurigobius friesii, Macroramphosus scolopax, Synchiropus phaeton* | | |
| **B** |  | International Bottom Trawl Survey in the Mediterranean (MEDITS) [87] |
| **P/B** | $\log M=-0.0066-0.279\cdot\log L_{\infty}+0.6543\cdot\log k+0.4634\cdot\log T$  Mortality (z)= F+M | Empirical equation from [88]; L_∞_ and k parameter from [117,123,200,207] |
| **Q/B** | $\log Q/B=6.37-1.5045\cdot\log T^{'}-0.168\cdot\log W_{\infty}+0.1399\cdot Pf+0.2765\cdot HD$ | Empirical equation from [93]; *a* and *b* parameters from [117,123,198] |
| **D** |  | [136,159,208–210] |
| **Di** |  | [110,111] |
| 39. **Bathydemersal (deep sea) fish (BDD):** *Argentina sphyraena, Glossanodon leioglossus, Trachyrincus scabrus* | | |
| **B** |  | International Bottom Trawl Survey in the Mediterranean (MEDITS) [87] |
| **P/B** | $\log M=-0.0066-0.279\cdot\log L_{\infty}+0.6543\cdot\log k+0.4634\cdot\log T$  Mortality (z)= F+M | Empirical equation from [88]; L_∞_ and k parameter from [117,211,212] |
| **Q/B** | $\log Q/B=6.37-1.5045\cdot\log T^{'}-0.168\cdot\log W_{\infty}+0.1399\cdot Pf+0.2765\cdot HD$ | Empirical equation from [93]; *a* and *b* parameters from [117,123,155] |
| **D** |  | [159,213] |
| **Di** |  | [110,111] |
| 40. **Small-spotted catshark (SPC):** *Scyliorhinus canicula* | | |
| **B** |  | International Bottom Trawl Survey in the Mediterranean (MEDITS) [87] |
| **P/B** | $\log M=-0.0066-0.279\cdot\log L_{\infty}+0.6543\cdot\log k+0.4634\cdot\log T$  Mortality (z)= F+M | Empirical equation from [88]; L_∞_ and k parameter from [214] |
| **Q/B** | $\log Q/B=6.37-1.5045\cdot\log T^{'}-0.168\cdot\log W_{\infty}+0.1399\cdot Pf+0.2765\cdot HD$ | Empirical equation from [93]; *a* and *b* parameters from [123] |
| **D** |  | [159] |
| **C** |  | [110,111] |
| 41. **Blackmouth catshark (BKC):** *Galeus melastomus* | | |
| **B** |  | International Bottom Trawl Survey in the Mediterranean (MEDITS) [87] |
| **P/B** | $\log M=-0.0066-0.279\cdot\log L_{\infty}+0.6543\cdot\log k+0.4634\cdot\log T$  Mortality (z)= F+M | Empirical equation from [88]; L_∞_ and k parameter from [215] |
| **Q/B** | $\log Q/B=6.37-1.5045\cdot\log T^{'}-0.168\cdot\log W_{\infty}+0.1399\cdot Pf+0.2765\cdot HD$ | Empirical equation from [93]; *a* and *b* parameters from [123] |
| **D** |  | [216] |
| **C** |  | [110,111] |
| 42. **Other small demersal sharks (SDS):** *Chimaera monstrosa, Etmopterus spinax, Squatina acanthia****s*** | | |
| **B** |  | International Bottom Trawl Survey in the Mediterranean (MEDITS) [87] |
| **P/B** | $\log M=-0.0066-0.279\cdot\log L_{\infty}+0.6543\cdot\log k+0.4634\cdot\log T$  Mortality (z)= F+M | Empirical equation from [88]; L_∞_ and k parameter from [123] |
| **Q/B** | $\log Q/B=6.37-1.5045\cdot\log T^{'}-0.168\cdot\log W_{\infty}+0.1399\cdot Pf+0.2765\cdot HD$ | Empirical equation from [93]; *a* and *b* parameters from [123,217,218] |
| **D** |  | [98,216,218,219] |
| **C** |  | [110,111] |
| 43. **Rays and skates (RSK):** *Leucoraja naevus, Raja asterias, Raja clavata, Raja miraletus* | | |
| **B** |  | International Bottom Trawl Survey in the Mediterranean (MEDITS) [87] |
| **P/B** | $\log M=-0.0066-0.279\cdot\log L_{\infty}+0.6543\cdot\log k+0.4634\cdot\log T$  Mortality (z)= F+M | Empirical equation from [88]; L_∞_ and k parameter from [220,221] |
| **Q/B** | $\log Q/B=6.37-1.5045\cdot\log T^{'}-0.168\cdot\log W_{\infty}+0.1399\cdot Pf+0.2765\cdot HD$ | Empirical equation from [93]; *a* and *b* parameters from [123] |
| **D** |  | [98,222,223] |
| **C** |  | [110,111] |
| 44. **Torpedos (TOR):** *Torpedo marmorata, Torpedo nobiliana* | | |
| **B** |  | International Bottom Trawl Survey in the Mediterranean (MEDITS) [87] |
| **P/B** | $\log M=-0.0066-0.279\cdot\log L_{\infty}+0.6543\cdot\log k+0.4634\cdot\log T$  Mortality (z)= F+M | Empirical equation from [88]; L_∞_ and k parameter from [117,224] |
| **Q/B** | $\log Q/B=6.37-1.5045\cdot\log T^{'}-0.168\cdot\log W_{\infty}+0.1399\cdot Pf+0.2765\cdot HD$ | Empirical equation from [93]; *a* and *b* parameters from [117] |
| **D** |  | [98] |
| **C** |  | [110,111] |
| 45. **Coastal benthic cephalopods (CBC):** *Octopus vulgaris* | | |
| **B** |  | [225] and International Bottom Trawl Survey in the Mediterranean (MEDITS) [87] |
| **P/B** | Mortality (z)= F+M | [226] |
| **Q/B** | $Q/B=0.0683+0.0474(W)$ | [227,228] |
| **D** |  | [229,230] |
| **C** |  | [110,111] |
| 46. **Benthopelagic cephalopods (BPC):** *Alloteuthis subulata, Illex coindetii, Todarodes sagittatus* | | |
| **B** |  | [231] and International Bottom Trawl Survey in the Mediterranean (MEDITS) [87] |
| **P/B** | Mortality (z)= F+M | [226] |
| **Q/B** | $Q/B=0.0683+0.0474(W)$ | [227,228] |
| **D** |  | [223,232–236] |
| **C** |  | [110,111] |
| 47. **Mesopelagic cephalopods (MEC):** *Histioteuthis reversa, Histioteuthis spp., Onychoteuthis banksii* | | |
| **B** |  | [231] and International Bottom Trawl Survey in the Mediterranean (MEDITS) [87] |
| **P/B** | Mortality (z)= F+M | [226] |
| **Q/B** | $Q/B=0.0683+0.0474(W)$ | [227,228] |
| **D** |  | [223] |
| **C** |  | [110,111] |
| 48. **Other benthic cephalopods (BCE):** *Bathypolypus sponsalis, Eledone cirrhosa, Octopus salutii, Pteroctopus tetracirrhus, Rossia macrosoma, Sepia elegans, Sepia orbignyana, Sepia spp, Sepietta oweniana, Sepiola robusta, Sepiola spp.* | | |
| **B** |  | [225] and International Bottom Trawl Survey in the Mediterranean (MEDITS) [87] |
| **P/B** | Mortality (z)= F+M | [226] |
| **Q/B** | $Q/B=0.0683+0.0474(W)$ | [227,228] |
| **D** |  | [223,229] |
| **C** |  | [110,111] |
| 49. **Bivalves (BIV):** *Bivalvia spp* | | |
| **B** |  | [237,238] |
| **P/B** |  | [225] |
| **Q/B** |  | [225] |
| **D** |  | [239] |
| **C** |  | [110,111] |
| 50. **Gastropods (GAS):** *Gastropoda spp* | | |
| **B** |  | [237,238] |
| **P/B** |  | [225] |
| **Q/B** |  | [240] |
| **D** |  | [241] |
| **C** |  | [110,111] |
| 51. **Deep-water rose shrimp (DWS):** *Parapenaeus longirostris* | | |
| **B** |  | International Bottom Trawl Survey in the Mediterranean (MEDITS) [87] |
| **P/B** | Mortality (z)= F+M | [226] |
| **Q/B** |  | [240] |
| **D** |  | [242] |
| **C** |  | [110,111] |
| 52. **Blue and red shrimp (BRS):** *Aristeus antennatus* | | |
| **B** |  | International Bottom Trawl Survey in the Mediterranean (MEDITS) [87] |
| **P/B** | Mortality (z)= F+M | [226] |
| **Q/B** |  | [240] |
| **D** |  | [243] |
| **C** |  | [110,111] |
| 53. **Giant red shrimp (GRS):** *Aristaeomorpha foliacea* | | |
| **B** |  | International Bottom Trawl Survey in the Mediterranean (MEDITS) [87] |
| **P/B** | Mortality (z)= F+M | [226] |
| **Q/B** |  | [240] |
| **D** |  | [242] |
| **C** |  | [110,111] |
| 54. **No commercial shrimps (NCS):** *Aegaeon lacazei, Calocaris macandreae, Chlorotocus crassicornis, Pasiphaea multidentata, Pasiphaea sivado, Philocheras echinulatus, Plesionika acanthonotus, Plesionika antigai, Plesionika edwardsii, Plesionika gigliolii, Plesionika heterocarpus, Plesionika martia, Plesionika spp, Pontophilus spinosus, Sergestes arachnipodus, Sergia robusta, Solenacera membranacea* | | |
| **B** |  | [225] |
| **P/B** | Mortality (z)= F+M | [226] |
| **Q/B** |  | [240] |
| **D** |  | [242,244–246] |
| **Di** |  | [110,111] |
| 55. **Norway lobster (NWL):** *Nephrops norvegicus* | | |
| **B** |  | International Bottom Trawl Survey in the Mediterranean (MEDITS) [87] |
| **P/B** | Mortality (z)= F+M | [226] |
| **Q/B** |  | [240] |
| **D** |  | [247] |
| **C** |  | [110,111] |
| 56. **European lobster (LOB):** *Palinurus elephas* | | |
| **B** |  | International Bottom Trawl Survey in the Mediterranean (MEDITS) [87] |
| **P/B** | Mortality (z)= F+M | [226] |
| **Q/B** |  | [240] |
| **D** |  | [248] |
| **C** |  | [110,111] |
| 57. **Other commercial decapods (OCD):** *Palinurus mauritanicus, Squilla mantis* | | |
| **B** |  | International Bottom Trawl Survey in the Mediterranean (MEDITS) [87] |
| **P/B** | Mortality (z)= F+M | [226] |
| **Q/B** |  | [240] |
| **D** |  | [249] |
| **C** |  | [110,111] |
| 58. **Non-commercial decapods (NDC):** *Dardanus arrosor, Inachus dorsettensis, Latreillia elegans, Liocarcinus corrugatus, Macropodia linaresi, Macropodia longipes, Macropodia spp, Macropipus tuberculatus, Monodaeus couchii, Munida intermedia, Munida rugosa, Munida spp., Pagurus excavatus, Pagurus prideaus, Paramola cuvieri, Polycheles typhlops* | | |
| **B** |  | [225] |
| **P/B** | Mortality (z)= F+M | [226] |
| **Q/B** |  | [240] |
| **D** |  | [244,250,251] |
| **Di** |  | [110,111] |
| 59. **Purple sea urchin (PSU):** *Paracentrotus lividus* | | |
| **B** |  | [252] |
| **P/B** | Mortality (z)= F+M | [226] |
| **Q/B** |  | [253] |
| **D** |  | [254] |
| **C** |  | [110,111] |
| 60. **Other sea urchins (OSU):** *Brissopsis lyrifera, Cidaris cidaris, Gracilechinus acutus, Spatangus purpureus* | | |
| **B** |  | [237,238] |
| **P/B** | Mortality (z)= F+M | [226] |
| **Q/B** |  | [253] |
| **D** |  | [255] |
| **C** |  | [110,111] |
| 61. **Sea cucumbers (SCU):** *Holothuria spp* | | |
| **B** |  | [237,238] |
| **P/B** | Mortality (z)= F+M | [226] |
| **Q/B** |  | [253] |
| **D** |  | [256] |
| **C** |  | [110,111] |
| 62. **Other macro-benthos (OMB):** *Asteroidea spp, Cnidaria spp., Ophiuroidea spp, Sipunculus spp, Porifera spp, Polychaeta spp, Nemertea spp* | | |
| **B** |  | [237,238,257] |
| **P/B** | Mortality (z)= F+M | [258,259] |
| **Q/B** |  | [258,259] |
| **D** |  | [255,260] |
| **Di** |  | [110,111] |
| 63. **Jellyfish (JLL):** *Aequorea forskalea, Aurelia aurita, Chrysaora hysoscella, Hydrozoa, Pelagia noctiluca, Pleurobrachia pileus, Rhizostoma pulmo* | | |
| **B** |  | International Bottom Trawl Survey in the Mediterranean (MEDITS) [87] |
| **P/B** |  | [226] |
| **Q/B** |  | [240] |
| **D** |  | [261] |
| 64. **Salps (SAL):** *Thalia democratic, Salpa fusiformis, Salpa maxima, Pyrosoma, Pyrosoma atlanticum* | | |
| **B** |  | [262] and International Bottom Trawl Survey in the Mediterranean (MEDITS) [87] |
| **P/B** |  | [226] |
| **Q/B** |  | [240] |
| **D** |  | [262] |
| 65. **Other corals and gorgonians (CGO):** *Bryozoa, Myriapora truncata, Alcyonium acaule, Alcyonium palmatum, Alcyonium spp., Callogorgia verticillata, Caryophyllia smithii, Epizoanthus spp., Eunicella cavolini, Eunicella filiformis, Eunicella singularis, Eunicella spp., Eunicella verrucosa spp., Sertularella spp.* | | |
| **B** |  | [240] |
| **P/B** |  | [226] |
| **Q/B** |  | [240] |
| **D** |  | [260] |
| 66. **Macro zooplankton (MAZ):** | | |
| **B** |  | [263] |
| **P/B** |  | [240] |
| **Q/B** |  | [240] |
| **D** |  | [264] |
| 67. **Meso and micro zooplankton (MMZ):** | | |
| **B** |  | [263] |
| **P/B** |  | [240] |
| **Q/B** |  | [240] |
| **D** |  | [265] |
| 68. **Suprabenthos (SUB):** | | |
| **B** |  | [225] |
| **P/B** |  | [266] |
| **Q/B** |  | [240] |
| **D** |  | [267] |
| 69. **Small phytoplankton (SMP):** | | |
| **B** |  | [268] |
| **P/B** |  | [269] |
| 70. **Large phytoplankton (LPH):** | | |
| **B** |  | [268] |
| **P/B** |  | [269] |
| 71. **Detritus (DET)** | | |
| **B** | log_10_ D = -2.41 + 0.954 log_10_ PP + 0.863 log_10_ E | [270] |
| **72. Discards (DIS)** | | |
| **Di** |  | [110,111,271] |
| **73. Midwater trawlers - France (MTW)** | | |
| **C** |  | [110] |
| **74. Bottom trawlers – Spain (BTW)** | | |
| **C** |  | [111] |
| **75. Long Liners - Spain (LLI)** | | |
| **C** |  | [111] |
