## Supplemental Appendix D for "Current and potential contributions of the Gulf of Lion Fisheries Restricted Area to fisheries sustainability in the NW Mediterranean Sea"

Table D.1. Confidence intervals used to describe the uncertainty for functional group (FG) and each input parameter of the balanced Ecopath model.

| FG number | FG name | Biomass | P/B | Q/B | Diet | Catch |
| --- | --- | --- | --- | --- | --- | --- |
| 1 | Bottlenose dolphins | ±80 | ±50 | ±50 | ±30 |  |
| 2 | Striped dolphins | ±80 | ±50 | ±50 | ±30 |  |
| 3 | Short-beaked common dolphin | ±80 | ±50 | ±50 | ±30 |  |
| 4 | Fin whale | ±80 | ±50 | ±50 | ±30 |  |
| 5 | Deep sea-cetacean feeders marine mammals | ±80 | ±50 | ±50 | ±30 |  |
| 6 | Endangered seabirds | ±80 | ±50 | ±50 | ±30 |  |
| 7 | Loggerhead turtles | ±80 | ±50 | ±50 | ±30 |  |
| 8 | Pelagic sharks | ±80 | ±50 | ±50 | ±50 |  |
| 9 | Non-commercial large pelagic fishes | ±80 | ±50 | ±50 | ±10 |  |
| 10 | Bluefin tuna | ±80 | ±50 | ±50 | ±30 | ±50 |
| 11 | Swordfish | ±80 | ±50 | ±50 | ±30 | ±50 |
| 12 | Other large pelagic fishes | ±80 | ±50 | ±50 | ±30 | ±50 |
| 13 | Mackerels | ±80 | ±50 | ±50 | ±30 | ±50 |
| 14 | Horse mackerels | ±80 | ±50 | ±50 | ±30 | ±50 |
| 15 | Other medium pelagic fishes | ±80 | ±50 | ±50 | ±30 | ±50 |
| 16 | European sardine | ±30 | ±50 | ±50 | ±30 | ±50 |
| 17 | European anchovy | ±30 | ±50 | ±50 | ±50 | ±50 |
| 18 | Other small pelagic fish | ±80 | ±50 | ±50 | ±80 | ±50 |
| 19 | Benthopelagic fishes | ±10 | ±50 | ±50 | ±50 | ±50 |
| 20 | No commercial meso (bathy) pelagic fishes | ±10 | ±50 | ±50 | ±30 | ±50 |
| 21 | Anglerfish | ±10 | ±50 | ±50 | ±10 | ±50 |
| 22 | European conger | ±10 | ±50 | ±50 | ±30 | ±50 |
| 23 | European hake | ±10 | ±50 | ±50 | ±50 | ±50 |
| 24 | Other commercial large demersal fish | ±10 | ±50 | ±50 | ±30 | ±50 |
| 25 | Poor cod | ±10 | ±50 | ±50 | ±30 | ±50 |
| 26 | Blue whiting | ±10 | ±50 | ±50 | ±30 | ±50 |
| 27 | Common pandora | ±10 | ±50 | ±50 | ±50 | ±50 |
| 28 | Sparidae | ±10 | ±50 | ±50 | ±30 | ±50 |
| 29 | Common dentex | ±10 | ±50 | ±50 | ±10 | ±50 |
| 30 | Red scorpionfish | ±30 | ±80 | ±50 | ±10 | ±50 |
| 31 | Scorpaenidae | ±10 | ±50 | ±50 | ±10 | ±50 |
| 32 | Labridae and serranidae | ±10 | ±50 | ±50 | ±10 | ±50 |
| 33 | Flatfishes | ±10 | ±50 | ±50 | ±30 | ±50 |
| 34 | Other commercial medium demersal fish | ±10 | ±50 | ±50 | ±60 | ±50 |
| 35 | No commercial medium demersal fish | ±10 | ±50 | ±50 | ±50 | ±50 |
| 36 | Red mullet | ±10 | ±50 | ±50 | ±30 | ±50 |
| 37 | Surmullet | ±10 | ±50 | ±50 | ±10 | ±50 |
| 38 | No commercial small demersal fish | ±10 | ±50 | ±50 | ±50 |  |
| 39 | Bathydemersal (deep sea) fish | ±10 | ±50 | ±50 | ±30 | ±50 |
| 40 | Small-spotted catshark | ±10 | ±50 | ±50 | ±30 | ±50 |
| 41 | Blackmouth catshark | ±10 | ±50 | ±50 | ±30 | ±50 |
| 42 | Other small demersal sharks | ±10 | ±50 | ±50 | ±50 | ±50 |
| 43 | Rays and skates | ±10 | ±50 | ±50 | ±30 | ±50 |
| 44 | Torpedos | ±10 | ±50 | ±50 | ±50 |  |
| 45 | Coastal benthic cephalopods | ±10 | ±50 | ±50 | ±60 | ±50 |
| 46 | Benthopelagic cephalopods | ±80 | ±50 | ±50 | ±50 | ±50 |
| 47 | Mesopelagic cephalopods | ±80 | ±50 | ±50 | ±30 | ±50 |
| 48 | Other benthic cephalopods | ±80 | ±50 | ±50 | ±30 | ±50 |
| 49 | Bivalves | ±50 | ±50 | ±60 | ±60 | ±50 |
| 50 | Gastropods | ±50 | ±50 | ±80 | ±60 | ±50 |
| 51 | Deep-water rose shrimp | ±10 | ±50 | ±50 | ±30 | ±50 |
| 52 | Blue and red shrimp | ±10 | ±50 | ±80 | ±30 | ±50 |
| 53 | Giant red shrimp | ±10 | ±50 | ±80 | ±30 | ±50 |
| 54 | No commercial shrimps | ±80 | ±50 | ±80 | ±30 |  |
| 55 | Norway lobster | ±10 | ±50 | ±80 | ±60 |  |
| 56 | European lobster | ±10 | ±50 | ±80 | ±30 |  |
| 57 | Other commercial decapods | ±10 | ±50 | ±80 | ±50 |  |
| 58 | Non-commercial decapods | ±50 | ±50 | ±80 | ±30 |  |
| 59 | Purple sea urchin | ±80 | ±50 | ±80 | ±30 |  |
| 60 | Other sea urchins | ±50 | ±50 | ±80 | ±50 |  |
| 61 | Sea cucumbers | ±50 | ±50 | ±80 | ±80 |  |
| 62 | Other macro-benthos | ±50 | ±60 | ±80 | ±60 |  |
| 63 | Jellyfish | ±80 | ±50 | ±80 | ±30 |  |
| 64 | Salps | ±80 | ±50 | ±80 | ±30 |  |
| 65 | Other corals and gorgonians | ±80 | ±50 | ±80 | ±60 |  |
| 66 | Macro zooplankton | ±30 | ±50 | ±60 | ±60 |  |
| 67 | Meso and micro zooplankton | ±30 | ±50 | ±60 | ±50 |  |
| 68 | Suprabenthos | ±10 | ±40 | ±80 | ±60 |  |
| 69 | Small phytoplankton | ±80 | ±50 |  |  |  |
| 70 | Large phytoplankton | ±80 | ±50 |  |  |  |
| 71 | Detritus | ±80 |  |  |  |  |
| 72 | Discards | ±80 |  |  |  |  |

Table D.2. Reference points used to develop the Fishing at Maximum Sustainable Yield (Fmsy) simulations for the continental slope of the Eastern Gulf of Lions (CoSEGoL) FRA.

| **GSAs** | **Species** | **Scientific name** | **Fcurrent** | **F0.1** | **Source** | **F0.1/Fcurrent** | **Reduction**  **(%)** |
| --- | --- | --- | --- | --- | --- | --- | --- |
| 1, 5, 6 and 7 | European hake | *Merluccius merluccius* | 1.14 | 0.23 | STECF 2018 | 0.20 | 79.82 |
| 7 | Red mullet | *Mullus barbatus* | 1.30 | 0.64 | STECF 2018 | 0.49 | 50.77 |
| 6 | Norway lobster | *Nephrops norvegicus* | 0.44 | 0.12 | STECF 2018 | 0.27 | 72.73 |
| 9, 10 and 11 | Deep-water rose shrimp | *Parapenaeus longirostris* | 1.68 | 0.74 | STECF 2018 | 0.44 | 55.95 |
| 6 | Blue and red shrimp | *Aristeus antennatus* | 0.73 | 0.42 | STECF 2018 | 0.58 | 42.47 |
| 9, 10 and 11 | Giant red shrimp | *Aristaeomorpha foliacea* | 1.12 | 0.57 | STECF 2018 | 0.51 | 49.11 |
| 6 and 7 | European anchovy | *Engraulis encrasicolus* | 1.76 | 0.47 | SPELMED 2018 | 0.27 | 73.30 |
| 6 and 7 | European sardine | *Sardina pilchardus* | 1.71 | 0.68 | SPELMED 2018 | 0.40 | 60.23 |
