## Supplemental Appendix E for "Current and potential contributions of the Gulf of Lion Fisheries Restricted Area to fisheries sustainability in the NW Mediterranean Sea"

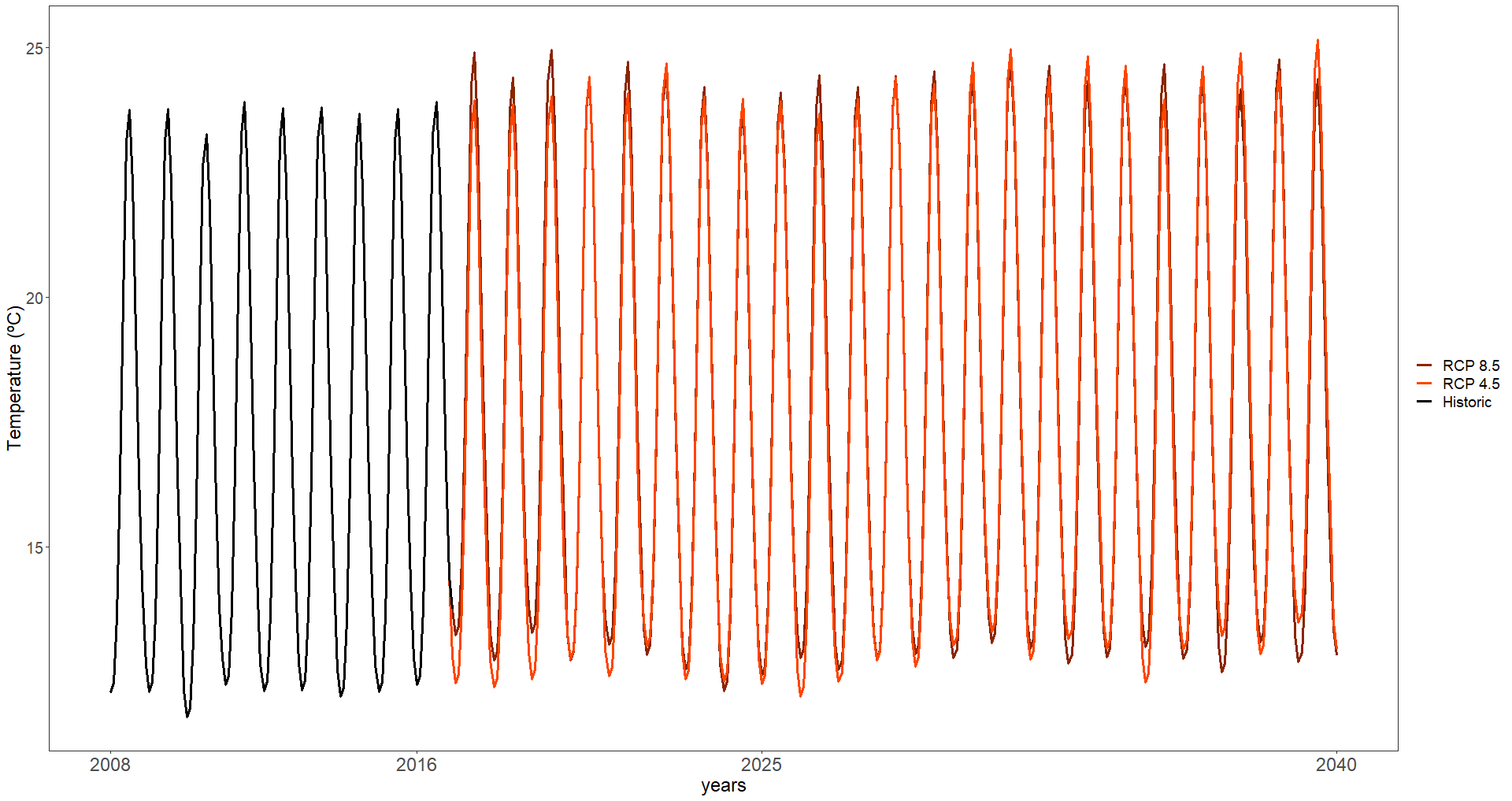

Figure E.1. Historic and future trends under the two scenarios of IPCC projections of sea water temperature in the continental slope of the Eastern Gulf of Lion Fisheries Restricted Area (CoSEGoL FRA) model.

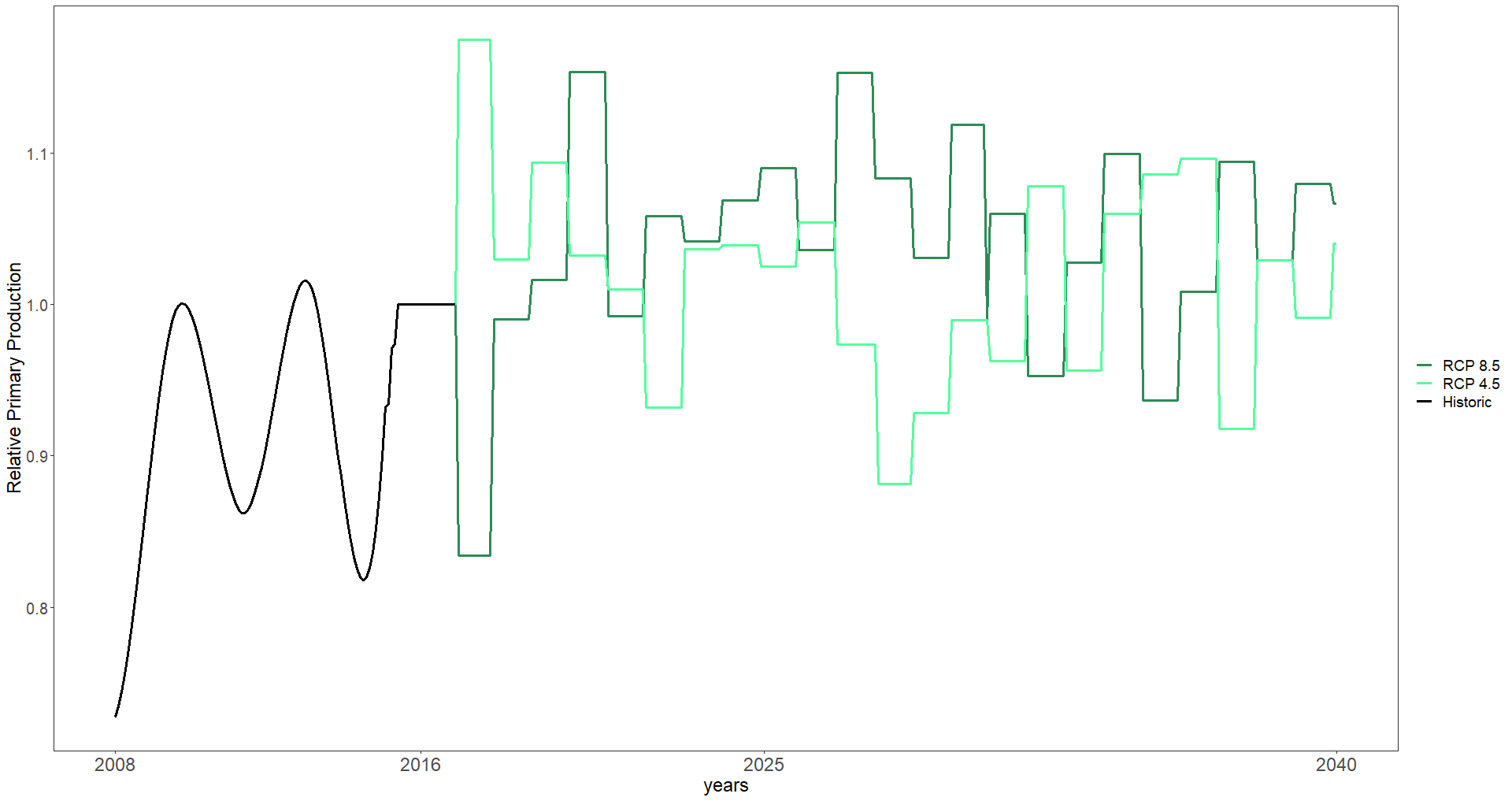

Figure E.2. Historic and future trends under the two scenarios of IPCC projections of relative primary production in the continental slope of the Eastern Gulf of Lion Fisheries Restricted Area (CoSEGoL FRA).
