## Supplemental Appendix F for "Current and potential contributions of the Gulf of Lion Fisheries Restricted Area to fisheries sustainability in the NW Mediterranean Sea"

Table F.1. Results of the fitting procedure of the CoSEGoL FRA ecosystem fitted to time series of data from 2008 to 2016. The table shows multiple fishing efforts to evaluate the sum of squares (SS) and AIC of each model. The best model chosen in this study is highlighted in bold.

| fishing effort (%) | vulnerabilities | spline points | SS | AIC | AICc |
| --- | --- | --- | --- | --- | --- |
| 10 | 10 | 2 | 360.18 | 157.84 | 160.53 |
| 10 | 10 | 4 | 355.74 | 161.65 | 165.45 |
| 10 | 10 | 6 | 353.24 | 166.28 | 171.39 |
| 10 | 20 | 2 | 337.11 | 178.81 | 189.31 |
| 10 | 20 | 4 | 333.92 | 184.20 | 197.04 |
| 10 | 20 | 6 | 316.21 | 184.92 | 200.40 |
| 10 | 30 | 2 | 310.05 | 205.10 | 230.53 |
| 10 | 30 | 4 | 300.32 | 209.80 | 239.32 |
| **10** | **30** | **6** | **289.45** | **214.39** | **248.44** |
| 5 | 10 | 2 | 335.33 | 149.90 | 152.60 |
| 5 | 10 | 4 | 327.57 | 152.50 | 156.29 |
| 5 | 10 | 6 | 308.04 | 151.09 | 156.19 |
| 5 | 20 | 2 | 318.89 | 172.64 | 183.14 |
| 5 | 20 | 4 | 313.82 | 177.32 | 190.15 |
| 5 | 20 | 6 | 309.71 | 182.61 | 198.09 |
| 5 | 30 | 2 | 304.09 | 202.94 | 228.38 |
| 5 | 30 | 4 | 297.35 | 208.69 | 238.22 |
| 5 | 30 | 6 | 286.46 | 213.24 | 247.29 |
| 1 | 10 | 2 | 369.49 | 160.67 | 163.37 |
| 1 | 10 | 4 | 360.67 | 163.18 | 166.97 |
| 1 | 10 | 6 | 347.46 | 164.45 | 169.56 |
| 1 | 20 | 2 | 353.09 | 183.95 | 194.45 |
| 1 | 20 | 4 | 349.08 | 189.13 | 201.97 |
| 1 | 20 | 6 | 338.91 | 192.61 | 208.09 |
| 1 | 30 | 2 | 329.81 | 211.95 | 237.39 |
| 1 | 30 | 4 | 315.31 | 215.20 | 244.73 |
| 1 | 30 | 6 | 300.83 | 218.67 | 252.72 |
| Baseline | 10 | 2 | 375.49 | 162.46 | 165.15 |
| Baseline | 10 | 4 | 358.11 | 162.39 | 166.18 |
| Baseline | 10 | 6 | 350.41 | 165.39 | 170.50 |
| Baseline | 20 | 2 | 357.39 | 185.29 | 195.79 |
| Baseline | 20 | 4 | 343.60 | 187.38 | 200.21 |
| Baseline | 20 | 6 | 332.89 | 190.62 | 206.10 |
| Baseline | 30 | 2 | 325.16 | 210.38 | 235.81 |
| Baseline | 30 | 4 | 316.17 | 215.51 | 245.03 |
| Baseline | 30 | 6 | 296.59 | 217.09 | 251.15 |
| -1 | 10 | 2 | 377.02 | 162.91 | 165.61 |
| -1 | 10 | 4 | 360.10 | 163.00 | 166.80 |
| -1 | 10 | 6 | 351.58 | 165.76 | 170.87 |
| -1 | 20 | 2 | 360.79 | 186.34 | 196.84 |
| -1 | 20 | 4 | 348.88 | 189.07 | 201.91 |
| -1 | 20 | 6 | 331.21 | 190.06 | 205.54 |
| -1 | 30 | 2 | 328.21 | 211.41 | 236.85 |
| -1 | 30 | 4 | 316.16 | 215.50 | 245.03 |
| -1 | 30 | 6 | 307.52 | 221.11 | 255.16 |
| -5 | 10 | 2 | 381.79 | 164.31 | 167.00 |
| -5 | 10 | 4 | 365.09 | 164.53 | 168.32 |
| -5 | 10 | 6 | 354.12 | 166.56 | 171.66 |
| -5 | 20 | 2 | 352.23 | 183.68 | 194.18 |
| -5 | 20 | 4 | 341.01 | 186.54 | 199.38 |
| -5 | 20 | 6 | 331.99 | 190.32 | 205.80 |
| -5 | 30 | 2 | 331.12 | 212.39 | 237.83 |
| -5 | 30 | 4 | 325.19 | 218.63 | 248.15 |
| -5 | 30 | 6 | 315.78 | 224.05 | 258.10 |
