## Supplemental Appendix A for "Current and potential contributions of the Gulf of Lion Fisheries Restricted Area to fisheries sustainability in the NW Mediterranean Sea"

The continental slope of the Eastern Gulf of Lions (CoSEGoL) Fisheries Restricted Area (FRA) model was developed using the Ecopath with Ecosim ecosystem modelling approach (EwE, version 6.6) and it was built using the best available information to represent the FRA ecosystem just before its establishment. Specifically, the model represented a situation of the CoSEGoL FRA for 2006-2008 time period. Ecopath is a mass-balanced model based on the assumption that the production of one functional group is equal to the sum of all predation, non-predatory losses, exports, biomass accumulations, and catches, as expressed by the following equation:

$$P/{B_{i}}\cdot B_{i}=P/{B_{i}}\cdot B_{i}\cdot\left( 1-{EE}_{i} \right)+\sum_{j} ({Q/B)}_{ji}\cdot B_{i}\cdot{DC}_{ji}+Y_{i}+{NM}_{i}+{BA}_{i}$$

where *B_i_* is the biomass, (*P/B)_i_* is the production rate, (*Q/B)_i_* is the consumption rate, *DC_ji_* is the fraction of prey *i* included in the diet of predator *j*, *NM_i_* is the net migration of prey *i*, *BA_i_* is the biomass accumulation of prey *i*, *Y_i_* is the catch of prey *i*, and *EE_i_* is the ecotrophic efficiency of prey *i*, that is, the proportion of production used in the system.

In an Ecopath model, the energy input and output of all functional groups must be balanced under some ecological and thermodynamic rules: (1) EE < 1.0; (2) P/Q [production/consumption rate or gross efficiency (GE)] ranges from 0.1 to 0.3 with the exception of fast growing groups such as bacteria; (3) R/A (respiration/food assimilation) < 1; (4) R/B (respiration/biomass) ranges from 1 to 10 for fishes and higher values for small organisms; (5) NE (net efficiency of food conversion	n) > GE and (6) P/R (production/respiration) < 1 [1,2]. To balance the FRA model, we applied a manual mass-balanced procedure following top-down approach modifying appropriate input parameters (starting from the functional groups with higher trophic level) and following the best practice guidelines provided in the literature [2,3].

Subsequently, an Ecosim model representing the CoSEGoL FRA ecosystem during the 2008−2016 period was fitted to time series of historical data. The Ecosim model describes the temporal dynamics of species biomass and flows over time by accounting for changes in predation, consumption rate, fishing and the environment [4,5]. Ecosim uses a set of differential equations to describe biomass dynamics:

$$\frac{dB_{i}}{dt}=\left( \frac{P}{Q} \right)_{i}\cdot\sum Q_{ji}-\sum Q_{ji}+I_{i}-\left( M_{i}+F_{i}-e_{i} \right)\cdot B_{i}$$

where *dB_i_/dt* is the growth rate of group *i* during time *t* in terms of its biomass *B_i_*; (P/Q)_i_ is the net growth efficiency of group *i*; *Q_ij_* is the consumption rate; *M_i_* is the non-predation mortality rate; *F_i_* is the fishing mortality rate; *e_i_* is the emigration; and *I_i_* is the immigration rate [4].

Consumption rates (*Q_ij_*) in Ecosim are calculated based on the “foraging arena” theory, which divides the biomass of the prey into a vulnerable and a non-vulnerable fraction and the transfer rate or vulnerability between the two fractions determines the trophic flow between the predator and the prey [6]. The vulnerability concept incorporates density-dependency processes and expresses how far a group is from its carrying capacity [1,4]. Default values of vulnerability (v = 2) represent a mixed trophic flow, a low value (v < 2) indicates ‘bottom-up’ flow control and a situation closer to carrying capacity, while a high value (v > 2) indicates ‘top-down’ flow control and a situation further away from carrying capacity [6,7]. For each predator-prey interaction, consumption rates are calculated as:

$Q_{ij}=\frac{a_{ij}\cdot v_{ij}\cdot B_{i}\cdot B_{j}\cdot T_{i}\cdot T_{j}\cdot{M_{ij}}/{D_{j}}}{v_{ij}+v_{ij}\cdot T_{i}\cdot M_{ij}+a_{ij}\cdot M_{ij}\cdot B_{j}\cdot{T_{j}}/{D_{j}}}\cdot f({Env}_{function},t)$,

where *a_ij_* is the rate of effective search for *i* by *j*; *T_i_* represents prey relative feeding time; *T_j_* the predator relative feeding time; *M_ij_* is the mediation forcing effects; *v_ij_* is the vulnerability parameter; *D_j_* represents the effects of handling time as a limit to consumption rate [1,6]; and *f(Env_function_,t)* is the environmental response function that restricts the size of the foraging arena (*C_rcj_*) to account for external environmental drivers changing over time, such as temperature [6,8].

The fitting procedure of the models was carried out with the Stepwise Fitting routine [9]. We fitted the Ecosim model to available time series of observed data for 2008-2016. The fitting allowed the estimation of prey-predator vulnerabilities and a primary production anomaly function based on minimizing the differences between predicted and observed data [4]. The primary production anomaly represents inter-annual variability of primary productivity, which often improve model fits [10]. The best model fits were selected based on optimized Akaike’s information criterion (AIC) for a maximum of 44 parameters in the CoSEGoL FRA. Finally, the best fits were manually chosen evaluating their credibility and sensibility of their behavior [2].
